# Translational and HIF1α-dependent metabolic reprograming underpin oncometabolome plasticity and synergy between oncogenic kinase inhibitors and biguanides

**DOI:** 10.1101/160879

**Authors:** Laura Hulea, Simon-Pierre Gravel, Masahiro Morita, Marie Cargnello, Oro Uchenunu, Young Kyuen Im, Shannon McLaughlan, Ola Larsson, Michael Ohh, Tiago Ferreira, Celia Greenwood, Gaëlle Bridon, Daina Avizonis, Josie Ursini-Siegel, Julie St-Pierre, Michael Pollak, Ivan Topisirovic

## Abstract

There is heightened interest to devise therapies that target the oncometabolome. We show that kinase inhibitors (KIs) and biguanides synergistically target melanoma, leukemia, and breast, colon and renal cancer cells, but not non-transformed cells. Metabolic profiling confirmed opposing effects of KIs and biguanides on glycolysis, but this was insufficient to explain the observed synergy between the drugs. Rather, we define a critical role for the synthesis of non-essential amino acids (NEAA) aspartate, asparagine and serine as well as reductive glutamine metabolism, in determining the sensitivity of cancer cells to KI - biguanide combinations. The mTORC1/4E-BP axis regulates aspartate, asparagine and serine synthesis by modulating translation of mRNAs encoding PC, ASNS, PHGDH and PSAT1. Ablation of 4E-BP1 and 2 results in a dramatic increase in serine, aspartate and asparagine levels and a substantial decrease in sensitivity of breast cancer and melanoma cells to KI - biguanide combinations. In turn, efficacy of KI – biguanide combinations is impeded by HIF1α and sustained reductive glutamine metabolism. These findings identify hitherto unappreciated translational reprograming of NEAA synthesis and HIF1α-dependent stimulation of reductive glutamine metabolism as critical metabolic vulnerabilities of cancer that underpin synergy between KIs and biguanides.

## Introduction

Metabolic perturbations are recognized as a hallmark of cancer (Hanahan and Weinberg, 2011). Metabolic reprograming is required to provide energy, building blocks and antioxidants to fuel neoplastic growth and protect cells from oxidative damage (Vander Heiden and DeBerardinis, 2017). It has been proposed that differences in metabolic programs between normal and cancer cells provide a therapeutic opportunity to target pathways that are essential for neoplastic growth, but largely dispensable in normal tissues (Vander Heiden, 2011). However, understanding of the targetable pathways that underlie metabolic vulnerabilities in cancer is incomplete. Aberrant activation of the proto-oncogenic kinases leads to changes in cell metabolism necessary for neoplastic growth (Pavlova and Thompson, 2016; Vander Heiden, 2011). Accordingly, the anti-neoplastic effects of clinically used kinase inhibitors (KIs) have been correlated to their ability to alter key metabolic pathways engaged by cancer cells, including glycolysis (Beloueche-Babari et al., 2015; Ko et al., 2016; Komurov et al., 2012). Metformin is a widely used anti-diabetic biguanide that induces energy stress by reducing oxidative phosphorylation via partial inhibition of complex I (Andrzejewski et al., 2014; Bridges et al., 2014; Wheaton et al., 2014). This leads to a compensatory increase in glucose uptake and glycolysis (Javeshghani et al., 2012). Biguanides exhibit anti-neoplastic effects in many cancer models (Anisimov et al., 2005; Ben Sahra et al., 2008; Cerezo et al., 2013; Dirat et al., 2015; Huang et al., 2008; Kisfalvi et al., 2009; Wheaton et al., 2014; Zakikhani et al., 2006). Metformin ameliorates type 2 diabetes at least in part by inhibiting gluconeogenesis, which leads to decreases in both glucose and insulin levels (Bailey and Turner, 1996; Wiernsperger and Bailey, 1999). These systemic effects of biguanides may contribute to their anti-neoplastic activity *in vivo* (Algire et al., 2011). However, there is compelling evidence that biguanides also inhibit tumor growth in a cell autonomous manner via direct induction of energy stress in cancer cells (Bridges et al., 2014; Chandel et al., 2016; Dowling et al., 2016; Fendt et al., 2013; Griss et al., 2015; Liu et al., 2016; Yuan et al., 2013). Phenformin is a more potent complex I inhibitor than metformin (Bridges et al., 2014). As phenformin use is associated with a higher risk of lactic acidosis than metformin (Assan et al., 1975), it is no longer used in diabetes treatment, but is nevertheless less toxic than many commonly used antineoplastic drugs.

Glycolysis is upregulated in most cancers, and is further increased by the inhibitors of oxidative phosphorylation (Gravel et al., 2014; Vander Heiden et al., 2009). Interventions that reduce glycolysis, including inhibitors such as 2-DG (Ben Sahra et al., 2010) or reduction in glucose concentration (Javeshghani et al., 2012) sensitizes neoplastic cells to biguanides. As KIs used in the clinic also reduce glycolysis (Pollak, 2013; Zhao et al., 2011), this is an attractive rationale for combining these drugs with biguanides. Indeed, biguanides potentiate effects of BRAF and ERK inhibitors in melanoma (Trousil et al., 2017; Yuan et al., 2013). However, the metabolic and signaling pathways that mediate anti-neoplastic effects of KI and biguanide combinations are largely unknown. Herein, oncometabolomes of cancer cells driven by diverse oncogenic kinases and different tissues of origin were systematically probed with combinations of KIs [lapatinib (HER2/EGFR inhibitor), PLX4032 (BRAF inhibitor) or imatinib (BCR-ABL inhibitor)] and phenformin. This revealed that metabolic reprograming via translational regulation of NEAA synthesis and HIF1α–dependent increase in reductive glutamine metabolism are key factors in determining the anti-neoplastic efficacy of KI – biguanide combinations and at the same time illuminated the plasticity of the oncometabolome.

## Results

### Phenformin and lapatinib exhibit synergistic anti-proliferative effect across different cancer cell lines

Combining KIs and phenformin has been shown to be effective in melanoma cells (Trousil et al., 2017; Yuan et al., 2013). We therefore first sought to establish the universality of these findings by combining clinically used KIs and phenformin in cancer cell lines of different origin that harbor diverse activated forms of oncogenic kinases. The effects of combination of phenformin and lapatinib on the proliferation were monitored in Normal Murine Mammary Gland (NMuMG) cells transformed with oncogenic Neu/ErbB2 (V664E) (rat homologue of HER2) (NMuMG–NT2197; hereafter referred to as NT2197), a well-established model of HER2-driven breast cancer (BCa) (Ursini-Siegel et al., 2008), lapatinib-responsive VHL-deficient renal cancer RCC4 cells (Brodaczewska et al., 2016) and HCT116 colorectal cancer (CRC) cells (Brattain et al., 1981). We also investigated the effects of phenformin and PLX4032 in A375 melanoma cells with mutated BRAF (Giard et al., 1973) and imatinib in K652 myelogenous leukemia cells which are BCR-ABL positive (Lozzio and Lozzio, 1975). Cells were treated with increasing concentrations of lapatinib, phenformin, or combination thereof for 72h and proliferation rates were assessed by BrdU incorporation. A concentration of phenformin (100μM) that only marginally inhibited proliferation was sufficient to dramatically sensitize NT2197 cells to lapatinib (Fig. 1A; Fig. S1A,D), whereas low concentrations of lapatinib (75nM) strongly increased anti-proliferative effects of phenformin (Fig. S1F,G). Notably, at these concentrations phenformin and lapatinib combination did not exert conspicuous effects on proliferation of non-malignant NMuMG cells (Fig. 1A; Fig. S1B,E). We next determined the nature of lapatinib and phenformin interaction in NT2197 cells by measuring combination index [CI50; (Berenbaum, 1989; Chou and Talalay, 1984; Tallarida, 2006)], which revealed that lapatinib and phenformin exhibit synergistic anti-proliferative effect (CI50 ∼0.79; Fig. 1B). Moreover, when combined, these drugs decreased survival of NT2197 cells to a much higher extent than either drug alone (59% of cells showed cumulated early and late apoptosis for dual phenformin-lapatinib treatment, compared to 16% and 13% for individual phenformin and lapatinib treatment, respectively; Fig 1C). Parallel results were observed in all the cell lines that were tested thus indicating that KIs and phenformin exhibit synergistic anti-neoplastic effects that are largely independent of the type of the driving oncogene and/or cancer origin (Fig. 1D-F, Fig. S1H-P).

**Fig. 1:**
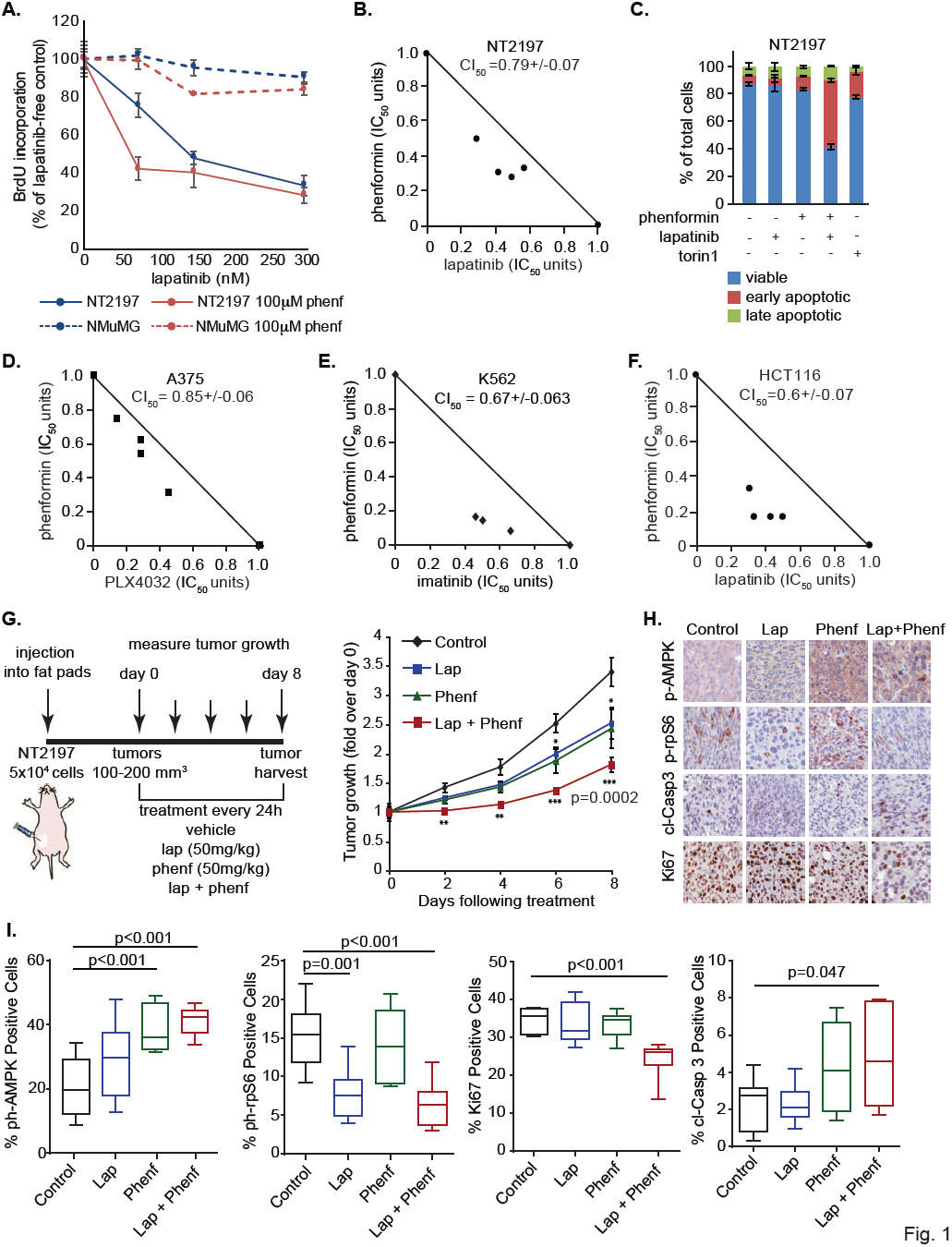
Kinase inhibitors and phenformin synergistically inhibit cell proliferation and restrict tumor growth *in vivo* A. Cell proliferation of NMuMG and NT2197 cells treated with the indicated concentrations of phenformin and lapatinib or combination thereof for 72h. Cell proliferation was measured by BrdU incorporation and expressed as percentage of non-lapatinib treated cells. The data are presented as mean values +/- the SD (n=3). **B.** The combined effect (CI_50_) of combination of lapatinib plus phenformin was evaluated in NT2197 cells by the isobologram method using the IC_50_ values and isobologram equation combination index (described in Material and Methods section). The CI_50_ individual values were generated from 3 independent experiments. **C.** NT2197 cells were treated with phenformin (250μM), lapatinib (600nM) or combination thereof, or torin1 (150nM) for 72h. Apoptosis was measured by AnnexinV-FITC and PI staining. The combined fractions (%) of early apoptotic (Annexin V+/ PI-) and late apoptotic (Annexin V+/PI+) cells are shown relative to the total cell population. The data are presented as mean values +/- the SD (n=3) and are representative of 3 independent experiments. **D, E, F.** A375 cells **(D)** were treated with phenformin and/or PLX4032 at different doses for 72h. K562 cells **(E)** were treated with phenformin and/or imatinib at different doses for 72h. HCT116 cells **(F)** were treated with phenformin and/or lapatinib at different doses for 72h. Cell proliferation was measured by BrdU incorporation. The combined effect of combination of PLX4032 **(D),** imatinib **(E)** or lapatinib (**F**) plus phenformin was evaluated by the isobologram method using the IC_50_ (as in **Fig. 1B**). The proliferation curves used to generate the isobolograms are shown in **Fig. S1H-M.** The CI_50_ individual values were generated from 2 independent experiments. **G-I**. 50,000 NT2197 cells were injected into two mammary fat pads of Nu/Nu mice. When tumors reached 100-200 mm^3^ mice were randomly distributed into 4 groups of 5 mice and treated with phenformin (50mg/kg), lapatinib (50mg/kg) or both every 48h for 8 days. **G**. Tumor growth was measured every two days until the control tumors reached 500mm^3^. The data are represented as mean values +/- the SEM (n=10) (\*\**P*< 0.001 and ***< 0.0005 (two-way ANOVA)). Mice were sacrificed on day 8, 4h after the last treatment. **I**. Tumor sections from 7 tumors/group were stained by immunohistochemistry using phospho-rpS6, phopsho-AMPK, Ki67 and cleaved Caspase3 (cl-Casp3) antibodies. (p-values determined by ANOVA). **H.** Representative images for staining quantified in **I.**

To establish whether, as reported for melanoma (Yuan et al., 2013), the effects observed in cell culture also occur *in vivo*, xenograft mammary tumors were generated by injecting NT2197 cells into the mammary fat pads of immunodeficient mice. Once the tumors reached an average size of 100-200 mm^3^, mice were treated with lapatinib (gavage; 50mg/kg), phenformin (IP; 50mg/kg), or a combination thereof and tumor growth was compared to control mice treated with a vehicle. While lapatinib or phenformin alone at these doses only modestly inhibited tumor growth (∼25% inhibition compared to control), their combination resulted in a significantly stronger anti-neoplastic effect (∼50% of control, p=0.0002) (Fig. 1G). Anti-tumorigenic effects of lapatinib were potentiated by phenformin as illustrated by a decrease in Ki67 (proliferation marker) and concomitant increase in the cleaved-caspase 3 (apoptosis marker) staining as compared to single lapatinib treatment (Fig. 1H,I). This shows that, similar to what was observed *in vitro*, the combined treatment has both a cytotoxic and cytostatic effect of tumor cells *in vivo*. Collectively, these results are consistent with previous observations in the melanoma model, as they demonstrate that phenformin increases anti-tumor efficacy of KIs *in vivo*.

### Lapatinib and biguanides have distinct impacts on the cancer cell metabolome

Lapatinib is a dual HER2/epidermal growth factor receptor (EGFR) tyrosine kinase inhibitor approved for treatment of BCa patients with amplified HER2 (Geyer et al., 2006). As HER2 engages a multitude of metabolic pathways (Jin et al., 2010; Walsh et al., 2013; Youngblood et al., 2016), we set out to identify potential metabolic vulnerabilities induced by lapatinib alone or by its combination with the biguanide phenformin. As expected, phenformin profoundly altered both glycolytic and citric acid cycle (CAC) intermediates levels (Fig. 2A, upper panel). Notably, phenformin increased lactate levels while strongly depleting citrate and succinate levels, supporting reduced CAC activity due to complex I inhibition and compensatory increase in glycolytic activity (Fendt et al., 2013; Javeshghani et al., 2012). On the contrary, lapatinib reduced lactate levels, in agreement with previous observation that KIs suppress glycolysis in cancer cell lines (Deblois et al., 2016; Pollak, 2013; Zhao et al., 2011), and reduce ^18^F-Fludeoxyglucose (FDG) uptake prior to any decrease in tumor volume in patients (Gebhart et al., 2013; McArthur et al., 2012). Strikingly, lapatinib and phenformin exhibited strong opposite effects on the CAC intermediates fumarate and malate, suggesting that KIs may impede metabolic adaptations induced in response to biguanides. Indeed, lapatinib attenuated the CAC intermediates changes induced by phenformin, as well as the amino acid aspartate (Fig.2A, lower panel). This effect was unique among all 16 amino acids analysed, which highlights a nodal role for aspartate together with the aforementioned CAC intermediates fumarate and malate. Additionally, the use of lapatinib and phenformin increased serine and glycine levels, while reducing methionine levels, which suggests that one-carbon metabolism may play a role in response to KI-biguanide combination.

**Fig. 2:**
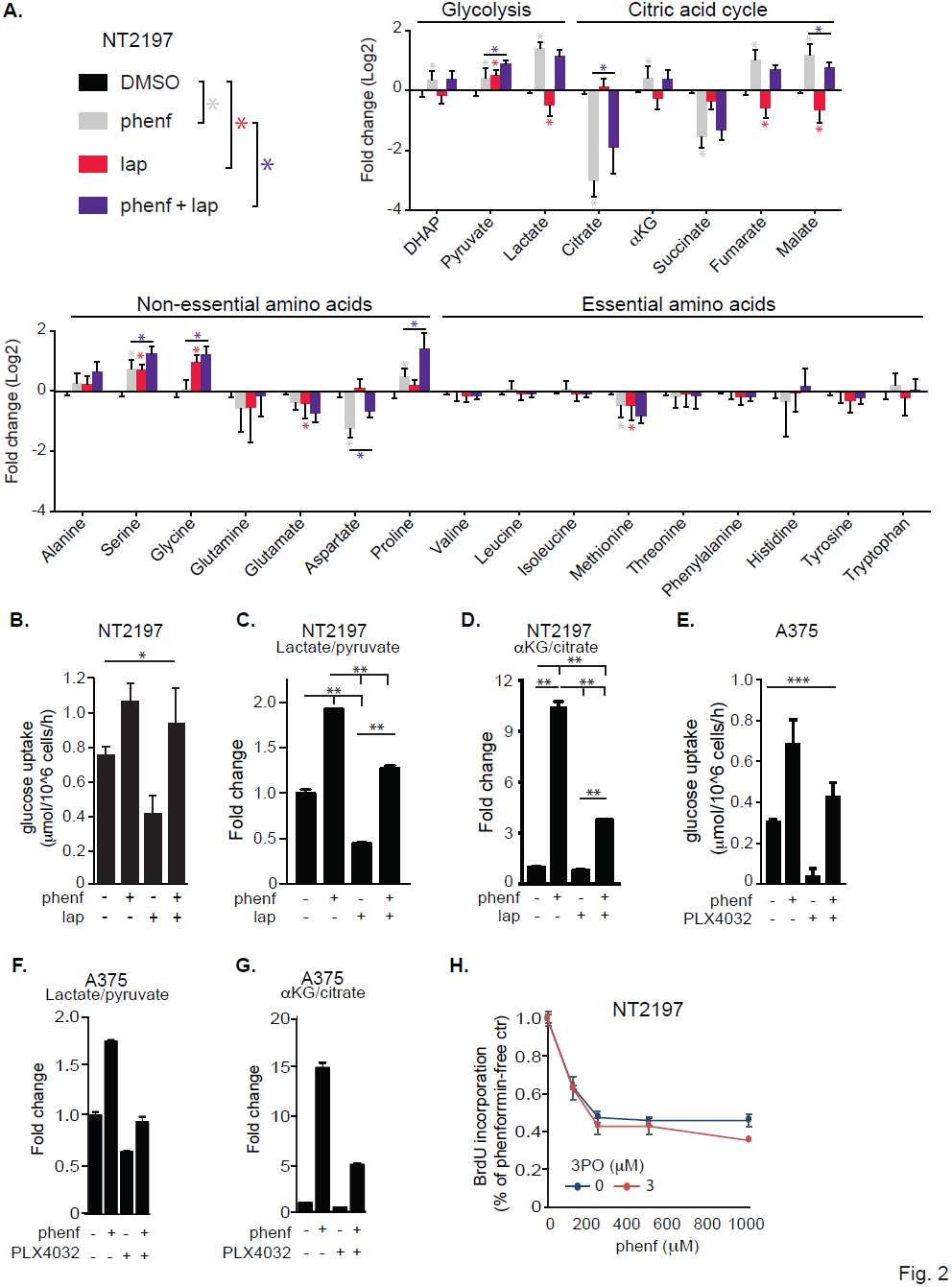
Phenformin blocks the metabolic adaptation induced by kinase inhibitors A. Levels of indicated metabolites in NT2197 cells treated for 24 hours with phenformin (600μM), lapatinib (600nM) or combination thereof were determined by GC–MS. Data are shown as mean +/- SD (3 independent experiments; \**P*< 0.05 (two-way ANOVA). **(B-D)** NT2197 cells were treated with phenformin (600μM) and/or lapatinib (600nM) for 24h. **B**. 24h post-treatment, glucose uptake was calculated. Data are shown as mean +/- SD (n=3; ** *P*< 0.001 (two-way ANOVA); representative of 3 independent experiments). **C, D**. Intracellular lactate to pyruvate ratio (**C**), which indicates lactate dehydrogenase activity (i.e. glycolysis), and α-ketoglutarate (αKG) to citrate ratio (**D**), indicator of glutamine-dependent reversal of citric acid cycle (reductive glutamine metabolism) were determined by GC/MS. Data are shown as mean +/- SEM (n=3; \**P*< 0.05 and **< 0.001 (two-way ANOVA); representative of 2 independent experiments). **E-G.** A375 cells were treated with phenformin (1.5mM) and/or PLX4032 (200nM) for 24h. **E**. 24h post-treatment, glucose uptake was calculated. Data are shown as mean +/- SD (n=3; \*\*\**P*< 0.0005 (two-way ANOVA); representative of 3 independent experiments). **F.** Intracellular lactate to pyruvate ratio, which indicates lactate dehydrogenase activity (i.e. glycolysis), was determined by GC/MS. Data are shown as mean +/- SEM (n = 3; representative of 2 independent experiments). **G.** α-ketoglutarate (αKG) to citrate ratio, indicator of glutamine-dependent reversal of citric acid cycle (reductive glutamine metabolism) was determined by GC/MS. Data are shown as mean +/- SEM (n=3; representative of 2 independent experiments).**H.** NT2197 cells were treated with the indicated concentrations of phenformin and 3PO or combination thereof for 72h. Cell proliferation was measured by BrdU incorporation and expressed as percentage of non-lapatinib treated cells. The data are presented as mean values +/- the SD (n=3; representative of 2 independent experiments).

As expected, glucose uptake and glycolysis were inhibited by lapatinib but increased by phenformin (Fig. 2B, C). Phenformin-induced increase in glycolysis (estimated by an increased lactate/pyruvate ratio) was significantly attenuated by lapatinib (Fig. 2C). Phenformin also increased α-ketoglutarate/citrate ratio which is indicative of elevated reductive glutamine metabolism (Fig. 2D) (Fendt et al., 2013). Reductive glutamine metabolism allows synthesis of CAC intermediates for biosynthetic pathways, including lipogenesis, under conditions wherein OXPHOS is inhibited, which leads to the impaired conversion of acetyl-CoA to citrate and subsequent increase in α-ketoglutarate/citrate ratio (Fendt et al., 2013; Gravel et al., 2014; Mullen et al., 2012). More recently, reductive glutamine metabolism was also implicated in maintaining REDOX homeostasis (Jiang et al., 2016). Lapatinib obliterated phenformin-induced increase in reductive glutamine metabolism (Fig. 2D). Parallel results were also observed in other cell lines (Fig. 2E-G, Fig. S2A). These data suggest that KIs hinder metabolic adaptations to energy stress induced by biguanides at least in part by reducing glycolysis and reductive glutamine metabolism.

### Antagonistic effects of lapatinib and phenformin on glucose uptake are insufficient to explain their synergy

Suppression of the compensatory increase in glucose uptake and glycolysis induced via inhibition of OXPHOS by biguanides has been reported to result in cell death (Ben Sahra et al., 2010). It is therefore plausible that the synergistic anti-proliferative effects between lapatinib and phenformin are a consequence of lapatinib-induced suppression of the adaptive increase in glucose uptake and glycolysis in response to phenformin (Fig. 2B,C,E,F). To assess the extent to which alterations in glycolysis underpin the observed synergy between lapatinib and phenformin, we investigated the effects of the combination of phenformin and the glycolytic inhibitor (3PO), which inhibits 6-phosphofructo-2-kinase/fructose-2,6-bisphosphatase 3 (PFKFB3) (Clem et al., 2008). 3PO was used instead of the more potent inhibitors of glycolysis (e.g. 2-DG), as this allowed inhibition of glucose uptake to a comparable extent to that observed in lapatinib treated cells (∼30%; Fig. 2B, Fig. S2C). Strikingly, although we selected a concentration of 3PO that inhibited glucose uptake to a level comparable to lapatinib (Fig. 2B, Fig. S2C), only lapatinib exhibited synergistic anti-proliferative effect with phenformin (Fig. 2H). This suggests that the inhibitory effect of lapatinib on glucose uptake is insufficient to explain its synergistic effect with phenformin (Fig. 2H; Fig. S2B,C).

### KIs and biguanides suppress protein synthesis and cooperatively downregulate mTORC1 signaling

Oncogenic kinases increase global protein synthesis and lead to reprograming of mRNA translation (Bhat et al., 2015). As mRNA translation is one of the most energy consuming processes in the cell (Buttgereit and Brand, 1995), to maintain energy homeostasis cancer cells must balance ATP production with the rates of protein synthesis (Morita et al., 2013). Suppression of protein synthesis is therefore a major step in adaptation to energetic stress, as sustained translation activity in energy-deficient cells leads to energy crisis and cell death (Leprivier et al., 2013). To determine whether the inability of the cells to reduce protein synthesis under energy stress contributes to the synergy between lapatinib and phenformin, polysomes, monosomes and ribosomal subunits were separated on sucrose gradients by ultracentrifugation (Gandin et al., 2014). The number of ribosomes engaged in polysomes is proportional to the translation activity in the cell (Warner et al., 1963). Phenformin (250μ M) suppressed mRNA translation to a higher extent than the lapatinib (600nM) as illustrated by reduction in polysome/monosome ratio (Fig. 3A). The drug combination resulted only in a marginal additional reduction in mRNA translation as compared to phenformin alone (Fig. 3A). These results exclude the possibility that the synergy between lapatinib and phenformin is caused by energy depletion induced by sustained protein synthesis when ATP production is limited.

**Fig. 3:**
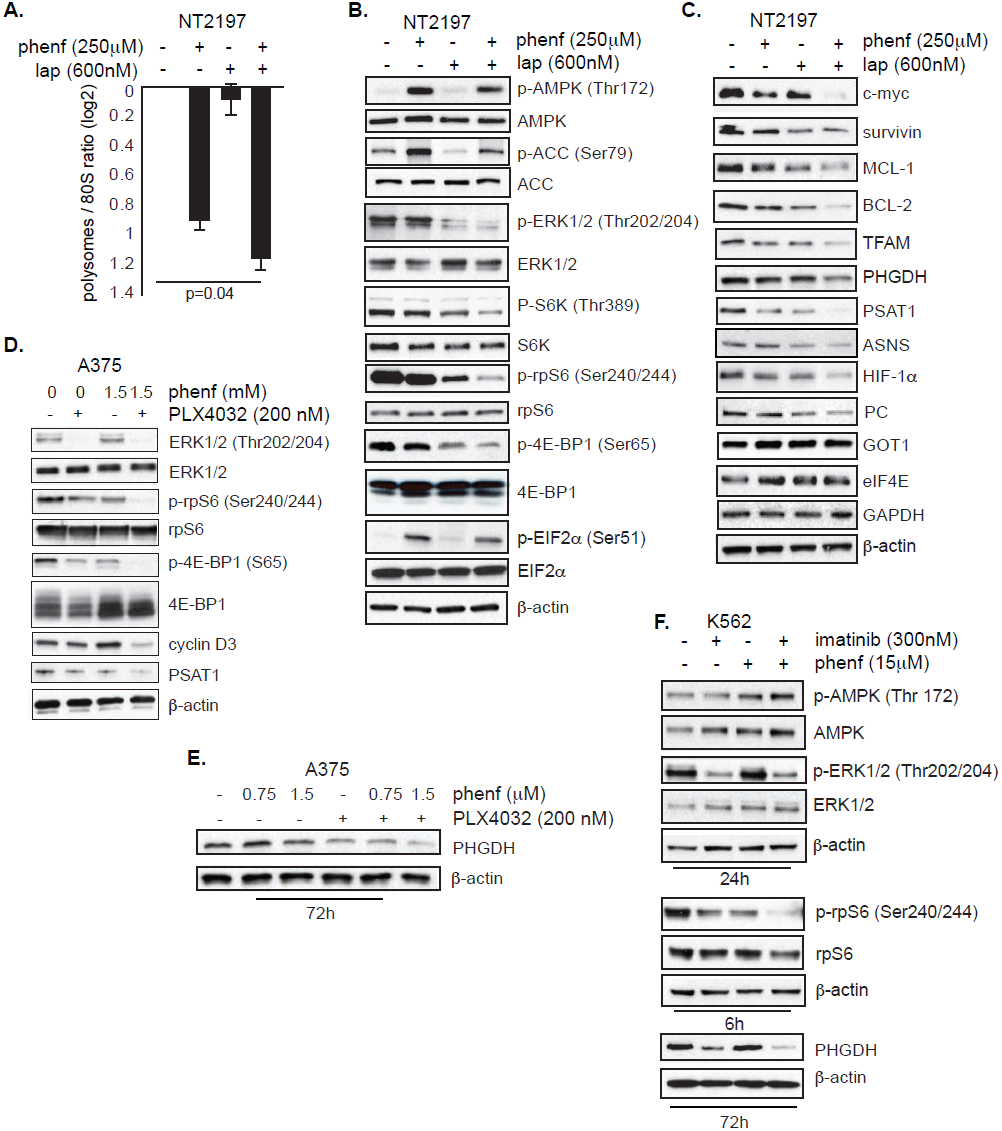
Kinase inhibitors and phenformin collaboratively inhibit mTORC1 A. NT2197 cells were treated with phenformin (250μM), lapatinib (600nM) or combination thereof for 4h, followed by polysome fractionation. Polysome to 80S ratios were calculated by comparing the area under the 80S peak and the combined area under the polysome (> 3 monosomes) peaks. Data from 2 independent experiments were used. Data were normalized to DMSO control, and ratios transformed into log2 base. The data are presented as mean values +/- the SEM (n=2 independent experiments; p-values determined by two-way ANOVA). **B**. NT2197 cells were treated with phenformin (250μM), lapatinib (600nM) or combination thereof for 6h. Levels of designated proteins were determined using western blot with indicated antibodies. β-actin served as a loading control. **C.** NMuMG-NT2197 cells were treated with phenformin (250μM), lapatinib (600nM) or combination thereof. Levels of designated metabolic proteins were determined using western blot with indicated antibodies. β-actin served as a loading control. **D, E.** A375 cells were treated as indicated for 6h **(D)** or 72h **(E)**. Expression and phosphorylation status of indicated proteins was determined by western blot using appropriate antibodies. β-actin served as a loading control. **F**. K562 cells were treated as indicated. Expression and phosphorylation status of indicated proteins was determined by western blot using appropriate antibodies. β-actin served as a loading control.

We next sought to determine the mechanisms of suppression of mRNA translation by lapatinib and phenformin combination. The mechanistic/mammalian target of rapamycin complex 1 (mTORC1) stimulates protein synthesis (Roux and Topisirovic, 2012). Lapatinib has been shown to inhibit mTORC1, and, in addition, persistent mTORC1 activity has been linked to the development of resistance to KIs (Lovly and Shaw, 2014). mTORC1 activity is also decreased by biguanides via the AMP-activated protein kinase (AMPK)-dependent and independent mechanisms (Ben Sahra et al., 2011; Dowling et al., 2007; Kalender et al., 2010). We therefore determined the effects of lapatinib, phenformin or combination thereof on mTORC1 signaling. Lapatinib (600nM) abolished the MAPK/ERK signaling in NT2197 cells as evidenced by reduction in ERK1/2 phosphorylation relative to the control, and suppressed mTORC1 activity as illustrated by decreased phosphorylation of two of its major substrates, the eukaryotic translation initiation factor 4E-binding protein 1 (4E-BP1) and the ribosomal protein S6 kinases 1 and 2 (S6K1/2), as well as ribosomal protein S6 (rpS6), which is a downstream target of S6K1/2 (Fig. 2B) (Grove et al., 1991; Kozma et al., 1990). As expected, phenformin (250μM) induced AMPK and phosphorylation of its downstream substrate acetyl-CoA carboxylase (ACC) (Fig. 3B), and suppressed mTORC1 (Fig. 3B). When the drugs were combined at the concentration which result in synergistic effect, mTORC1 was suppressed to a higher extent as compared to either drug alone (Fig. 3B). In addition, mTORC1 inhibition engendered by combining phenformin with PLX4032 in A375 cells or imatinib in K562 cells was more pronounced that when individual treatments were employed (Fig. 3D-F).

Intriguingly, the inhibition of mTORC1 by the combination of phenformin and lapatinib at concentrations that resulted in synergistic anti-proliferative effect, was less than that observed using the active site mTOR inhibitor, torin1 (Fig. S3B). This moderate mTORC1 inhibition achieved by the phenformin and lapatinib combination resulted in a pronounced pro-apoptotic effect (∼59% of cumulative early and late apoptosis cells; Fig. 1C), whereas almost total obliteration of mTORC1 signaling with torin1 lead to mostly cytostatic effect (∼23% of cumulative early and late apoptosis cells; Fig. 1C). This suggests that the synergy between phenformin and lapatinib is not attributable to more complete mTORC1 inhibition, but rather is mediated by moderate cooperative suppression of mTORC1 in concert with the effects of the drugs on glycolysis and reductive glutamine metabolism.

In addition to mTORC1, eIF2α phosphorylation, which limits amounts of ternary complex that recruits initiator tRNA during translation initiation, is implicated in regulation of protein synthesis under stress (Ron D, 2007). While phenformin strongly stimulated eIF2α phosphorylation, lapatinib alone or in combination with phenformin did not affect phospho-eIF2α levels (Fig. 3B), which explains more pronounced inhibition of global protein synthesis by phenformin as compared to lapatinib at indicated concentration of the drugs (Fig. 3A). Thus, the modulation of eIF2α phosphorylation is unlikely to underpin the synergistic anti-proliferative effects of phenformin and lapatinib. Collectively, these data indicate that in addition to exhibiting opposing effects on glycolysis and reductive glutamine metabolism, lapatinib and phenformin cooperatively suppress mTORC1, which in addition to the induction of eIF2α phosphorylation by phenformin results in reduction in protein synthesis.

### 4E-BPs are essential for synergistic effects of KI and biguanides

In addition to regulating global protein synthesis, mTOR inhibition leads to selective perturbations in the translatome whereby translation of specific subsets of mRNAs appear to be more sensitive to changes in mTOR activity than the others (Bhat et al., 2015; Hsieh et al., 2012; Larsson et al., 2012; Thoreen et al., 2012). This is in a large part mediated by the inactivation of 4E-BPs, and consequent upregulation of the eIF4F complex levels (Brunn et al., 1997; Gingras et al., 1999; Gingras et al., 1998; Pause et al., 1994). Increase in eIF4F complex levels selectively stimulates synthesis of factors implicated in oncogenesis (e.g. BCL-2 family members, cyclins, c-MYC) and nuclear-encoded proteins with mitochondrial functions (e.g. TFAM, components of complex I, III and V) (Gandin et al., 2016; Morita et al., 2013; Roux and Topisirovic, 2012). We observed that the anti-proliferative and pro-apoptotic effects of the lapatinib and phenformin combination (Fig. 1A-C) were associated with reduction in levels of proteins encoded by mRNAs previously demonstrated to be sensitive to fluctuations in eIF4F levels, including anti-apoptotic (survivin, MCL-1, BCL-2), oncogenic (c-MYC) and factors with mitochondrial function (TFAM), but not of proteins whose expression is less influenced by eIF4F levels, such as β-actin, GAPDH or eIF4E (Fig. 3C) (De Benedetti et al., 1994; Gandin et al., 2016; Larsson et al., 2012; Morita et al., 2013). (Fig. 3C).

Since the expression of proteins encoded by mRNAs that are known to be regulated by the mTORC1/4E-BP/eIF4F axis was more dramatically decreased in the cells treated with the combination of the drugs than either drug alone, we next investigated whether this axis contributes to synergistic anti-neoplastic effects between phenformin and lapatinib. To this end, we generated NT2197 cells that lack both 4E-BP1 and 4E-BP2 expression using CRISPR/Cas9 technology (Fig. 4A). The loss of 4E-BP1/2 expression strongly attenuated the anti-proliferative and pro-apoptotic effects of lapatinib and phenformin combination as compared to 4E-BP1/2 proficient cells (Fig. 4B-D, Fig. S4B-G) or 4E-BP1/2 deficient cells in which 4E-BP1 was re-expressed (Fig. S4H,I). Moreover, the loss of 4E-BP1/2 expression obliterated the effects of the drug combination on the eIF4F complex assembly, as monitored by m^7^ GTP pull-down assay and proximity ligation assay (PLA) (Fig. 4E-F, Fig. S4I). This was paralleled by diminished effects of phenformin and lapatinib combination on the expression of proteins encoded by mRNAs that are “eIF4F-sensitive” including survivin, BCL-2, TFAM, protein levels (Fig. 4G). The loss of 4E-BP1/2 expression also attenuated the anti-proliferative effects of PLX4032 and phenformin combination in melanoma cell model (Fig. 4H). These findings show that 4E-BPs are critical for the synergy between lapatinib and phenformin.

**Fig. 4:**
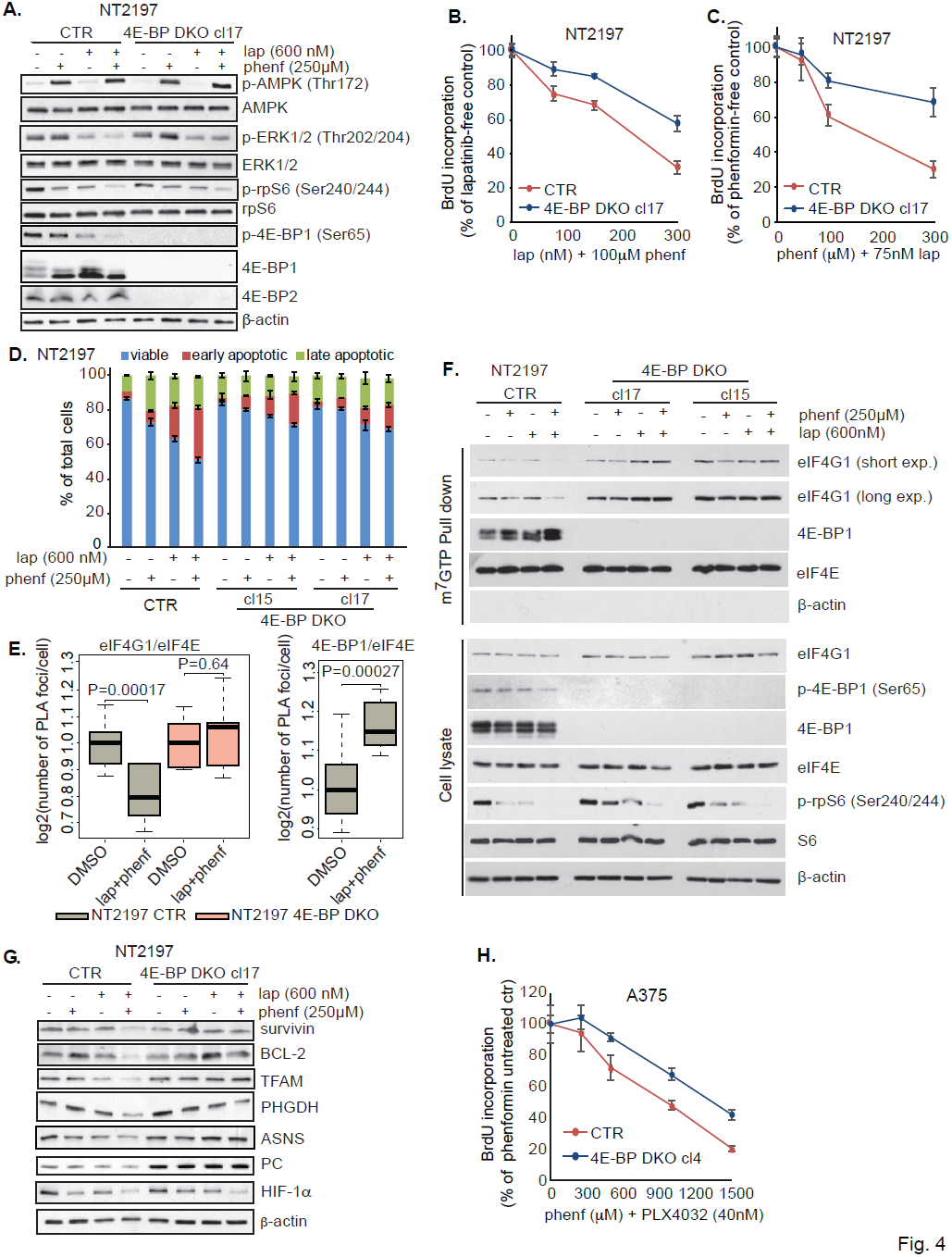
4E-BP1/2 mediate the response of NT2197 cells to the combination of lapatinib and phenformin A. NT2197 cells, control (CTR) or depleted of 4E-BP1 and 4E-BP2 by CRISPR (4E-BP DKO clone 17), were treated with phenformin (250μM) and lapatinib (600nM) or combination thereof for 6h. The expression and phosphorylation status of indicated proteins were determined by western blot using appropriate antibodies. β-actin served as a loading control. **B, C.** NT2197 cells, CTR or 4E-BP DKO cl17, were treated with the indicated concentrations of phenformin and lapatinib or combination thereof for 72h. Cell proliferation was measured by BrdU incorporation and expressed as percentage of non-lapatinib treated cells (**B**), or non-phenformin treated cells (**C**). The data are presented as mean values +/- the SD (n=3; representative of 3 independent experiments). **D.** NT2197 cells, CTR or 4E-BP DKO (cl15 and cl17), were treated with phenformin (250μM) and lapatinib (600nM) or combination thereof for 72h. Apoptosis was measured using a PE-Annexin V – 7-AAD staining and analyzed by flow cytometry. The fractions (%) of viable (Annexin V-/ 7-AAD-), early apoptotic (Annexin V+/ 7-AAD-) and late apoptotic (Annexin V+/7-AAD+) cells are shown relative to the total cell population. Results represent means +/- SD (n=3; representative of 3 independent experiments). **E.** NT2197 cells, CTR (grey) or 4E-BP DKO cl17 (red), were treated with the indicated phenformin (250μM) and lapatinib (600nM) or combination thereof for 4h. The interactions between eIF4E/eIF4G1 and eIF4E/4E-BP1 were assessed by proximity ligation assay. P-values were calculated using one-way ANOVA; posthoc Tukey test was applied. Representative images are shown in **Supplemental item 1. F**. NT2197 cells, CTR or 4E-BP DKO (cl15 and cl17), were treated with the indicated concentrations of phenformin and lapatinib or combination thereof for 4h. Protein lysates were subjected to m^7^GDP pull-down. Amounts of the indicated proteins in the input or pull-down were determined by western blotting; bb-actin served as a loading control (input) and to exclude contamination (m^7^GDP pull-down). **G**. NT2197 cells, CTR or 4E-BP DKO cl17, were treated with phenformin (250μM) and lapatinib (600nM) or combination thereof. The expression of indicated metabolic proteins were determined by western blot using appropriate antibodies. β-actin served as a loading control. **H**. A375 cells, control (CTR) or depleted of 4E-BP1 and 4E-BP2 by CRISPR (4E-BP DKO clone 4), were treated with the indicated concentrations of phenformin and plx4032 (37.5nM) for 72h. Cell proliferation was measured by BrdU incorporation and expressed as percentage of non-phenformin treated cells. The data are presented as mean values +/- the SD (n=3; representative of 3 independent experiments).

### The mTORC1/4E-BP axis regulates translation of mRNAs encoding rate-limiting enzymes of metabolic pathways that fuel neoplastic growth

Protein synthesis is one of the most energy-demanding cellular processes and therefore cancer cells, which have elevated levels of mRNA translation, must adapt their metabolism (Buttgereit and Brand, 1995; Morita et al., 2015). Emerging data suggest critical roles for serine aspartate and asparagine biosynthesis pathways in mTOR-dependent regulation of neoplastic growth and signaling (DeBerardinis and Chandel, 2016; Krall et al., 2016; Vander Heiden and DeBerardinis, 2017). Notably, the combination of phenformin and KIs decreased levels of proteins that play key roles in these metabolic pathways including phosphoglycerate dehydrogenase (PHGDH), phosphoserine aminotransferase 1 (PSAT1), pyruvate carboxylase (PC) and asparagine synthetase (ASNS), but not glutamic-oxaloacetic transaminase 1 (GOT1) in NT2197 (Fig. 3C), A375 and K562 cells (Fig. 3D-F; Fig. S4J) to a higher extent than each drug alone. Ablation of 4E-BP1 and 2, almost completely abolished the effects of phenformin and lapatinib on the expression of the aforementioned proteins in NT2197 cells (Fig. 4B).

We next assessed whether the effect of the lapatinib and phenformin combination on the expression of PHGDH, PSAT1, PC and ASNS occurs at the level of mRNA translation by using polysome profiling, which allows separation of efficiently (associated with heavy polysomes) vs. non-efficiently (associated with light polysomes) translated mRNAs based on the differences in their sedimentation in sucrose gradients (Gandin et al., 2014) (Fig. 5A). These experiments revealed that under basal conditions PHGDH, PSAT1, PC and ASNS mRNAs are translated at high efficiency as illustrated by the presence of these transcripts in the heavy polysome fractions (Fig. 5B). Lapatinib and phenformin combination reduced translation of PHGDH, PSAT1, PC and ASNS mRNAs in 4E-BP1/2 proficient, but not deficient cells as evidenced by their shift from the fractions containing more ribosomes to those with less ribosomes (Fig. 5B). A similar trend was observed for cyclin D3 mRNA, which is a previously established translational target of the mTORC1/4E-BP pathway (Larsson et al., 2012). In contrast, the lapatinib and phenformin combination exerted only a marginal effect on translation of housekeeping mRNAs (β-actin, GAPDH) or GOT1, which shows that the effects of the drug combination on translation of PHGDH, PSAT1, PC and ASNS mRNAs are selective (Fig. 5B). Collectively, these findings demonstrate that lapatinib and phenformin cooperatively reduce PHGDH, PSAT1, PC and ASNS protein levels in part by decreasing translational efficiency of corresponding mRNAs via suppression of the mTORC1/4E-BP axis. Importantly, depletion of ASNS and PHDGH in NT2197 cells lacking 4E-BP1/2 bolstered anti-proliferative effects of the lapatinib and phenformin combination (Fig. 6E,G). Changes in protein levels of PHGDH, PSAT1, and ASNS were not accompanied by the changes in mRNA levels, whereas PC mRNA was slightly increased in 4E-BP1/2 deficient vs. proficient cells (Fig. S5A,B). Collectively, these data show that the mTORC1/4E-BP-dependent translation control plays a major role in regulating levels of enzymes in serine, aspartate and asparagine biogenesis pathways.

**Fig. 5:**
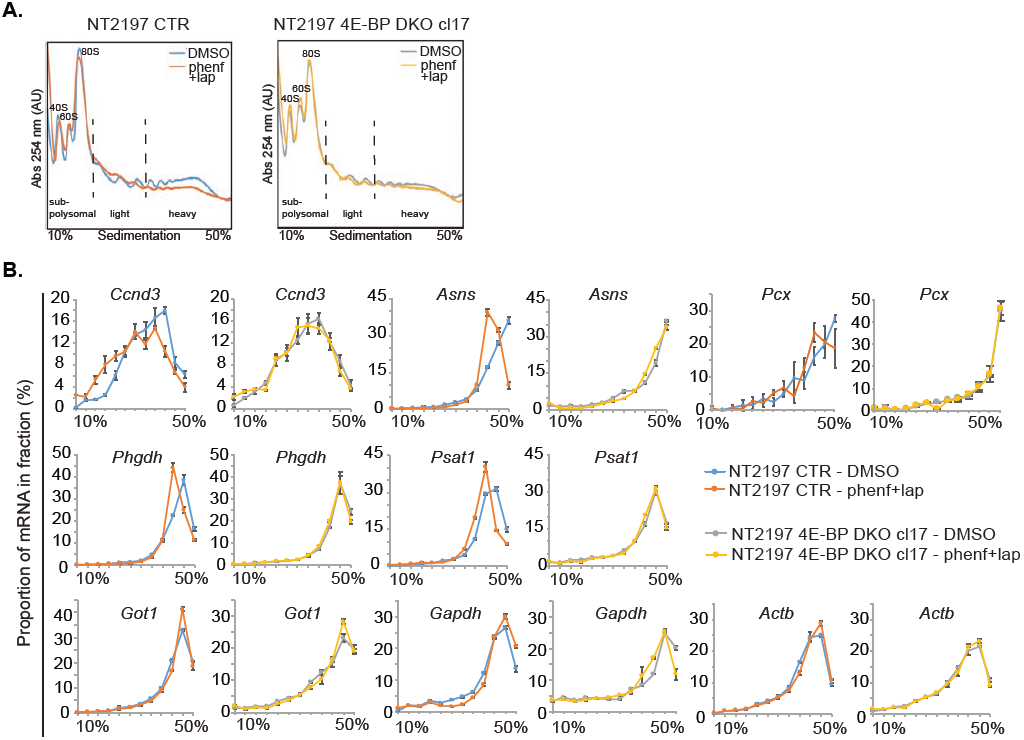
Phenformin disrupts the formation of the eIF4F complex A. NT2197 cells, CTR or 4E-BP DKO cl17, were treated with vehicle control or combination of phenformin (250μM) and lapatinib (600nM) for 4h. Sub-polysomal, light polysome and heavy polysome fractions were obtained from cytosolic extracts by ultracentrifugation using 5-50% sucrose gradients. During fractionation UV absorbance at 254nm (Abs 254nm) was continuously monitored to obtain absorbance tracings. Position of 40S and 60S ribosomal subunits, monosome (80S) and polysomes are indicated. **B.** Amount of indicated mRNAs in polysome fractions isolated from cells described in (**A**) were determined by RT-qPCR. Experiments in panels (**C**) and (**B**) were carried out in 3 independent experiments, each consisting of a triplicate. Data are representative of the 3 independent experiments and are expressed as a percentage of a given mRNA in each fraction +/- SD.

**Fig. 6:**
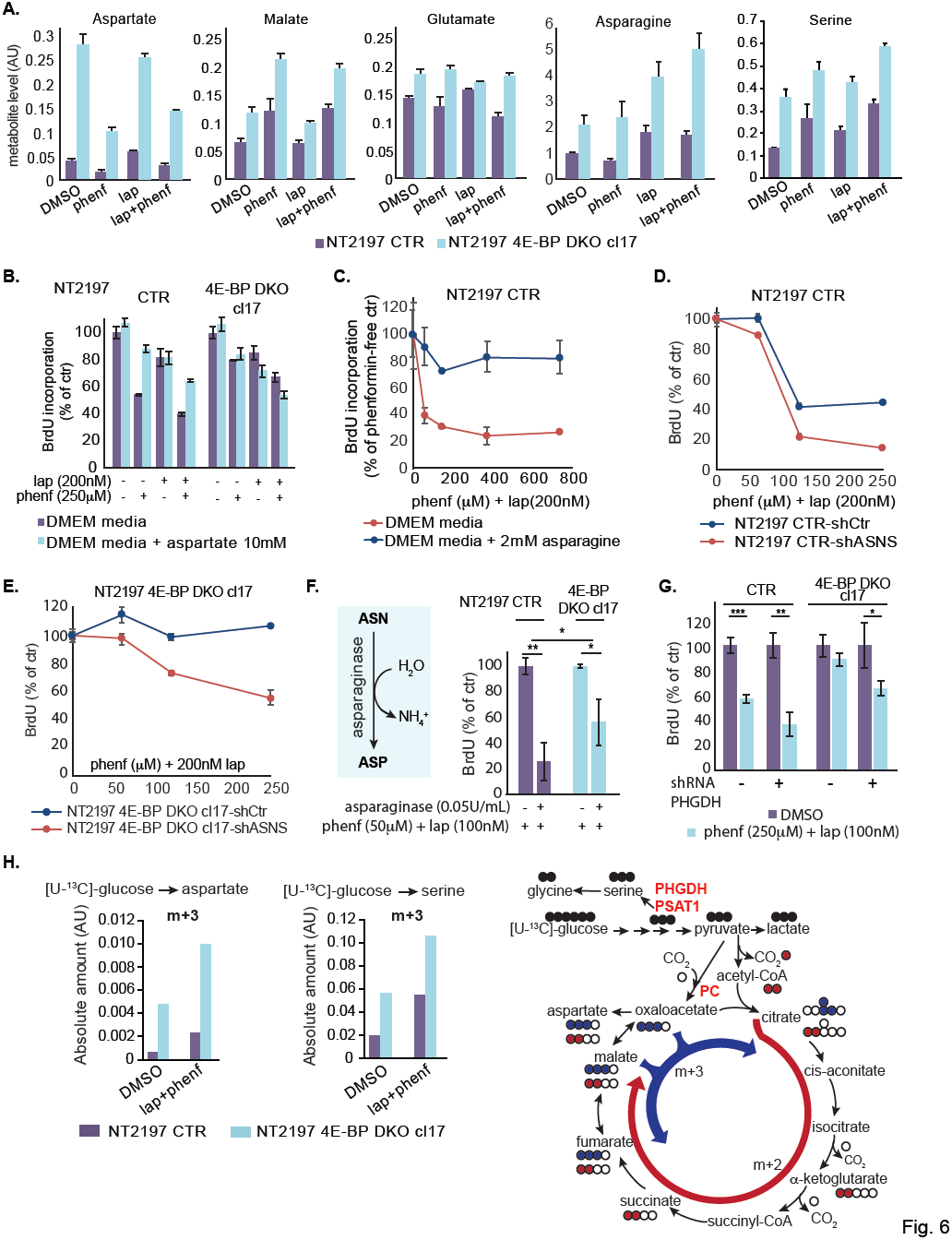
Metabolic reprograming of 4E-BP1/2 depleted cells decreases the efficacy of lapatinib and phenformin A. NT2197 cells, CTR or 4E-BP DKO cl17, were treated with phenformin (250μM) and lapatinib (600nM) or combination thereof for 24h. Levels of aspartate, malate, glutamate, serine and asparagine were determined by GC–MS. Data represent means +/- SEM (n=3; representative of 2 independent experiments). **B, C**. NT2197 cells, CTR or 4E-BP DKO cl17, were treated with the indicated concentrations of phenformin, lapatinib or combination thereof for 72h. Where indicated media were supplemented with aspartate (10mM) (**B**) or asparagine (2mM) (**C**). Cell proliferation was measured by BrdU incorporation and expressed as percentage of control treated cells. The data are presented as mean values +/- the SD (n=3; representative of 3 independent experiments). **D, E.** NT2197 cells, CTR or 4E-BP DKO cl17, expressing either control shRNA or ASNS shRNA, were treated with the indicated concentrations of phenformin and lapatinib for 72h. Cell proliferation was measured by BrdU incorporation and expressed as percentage of control treated cells. The data are presented as mean values +/- the SD (n=3; representative of 3 independent experiments). **F.** CTR or 4E-BP DKO cl17 NT2197 cells, were treated with the indicated concentrations of phenformin, lapatinib and asparaginase for 72h. Cell proliferation was measured by BrdU incorporation and expressed as percentage of control treated cells. The data are presented as mean values +/- the SD (n=3; \**P*< 0.05 and **< 0.001 (ANOVA); the test for interaction comparing the decrease in proliferation in CTR vs 4E-BP DKO cl17 after asparaginase treatment was significant (p=0.00803) and calculated from data from 2 indeepndent experiments). **G.** CTR or 4E-BP DKO cl17 NT2197 cells, expressing either control shRNA or PHGDH shRNA, were treated with the indicated concentrations of phenformin and lapatinib for 72h. Cell proliferation was measured by BrdU incorporation and expressed as percentage of control treated cells. The data are presented as mean values +/- the SD (n=3; \**P*< 0.05, **p< 0.001 and ***p< 0.0001 (one-way ANOVA)). **H**. CTR or 4E-BP DKO cl17 NT2197 cells, treated with vehicle control or a combination of phenformin (250mM) and lapatinib (600nM) for 24h, were incubated with ^13^C_6_-glucose for various time points. Data shown correspond to 10min incubation. Stable isotope tracer analyses were performed and the levels of aspartate (m+3) and serine (m+3) ion amounts are shown, expressed as absolute amount. Data arerepresentative of 2 independent experiments. AU: arbitrary units. Right panel: schematic representation of the ^13^C atom incorporation into metabolites.

### Translationally-induced metabolic reprograming caused by 4E-BP1/2 loss decreases the efficacy of lapatinib and phenformin

Loss of 4E-BP1 and 2 expression abolished effects of the combination of lapatinib and phenformin on translation of mRNAs encoding proteins which play critical role in aspartate, asparagine and serine synthesis. Moreover, as shown previously, suppression of mTOR signaling and subsequent 4E-BP1/2 activation is paralleled by reduction in translation of mRNAs encoding proteins with mitochondrial function (e.g. TFAM) (Fig. 5B) (Morita et al., 2013). We therefore studied the effects of 4E-BP1/2 ablation on steady-state metabolite levels. The 4E-BP1/2 loss engendered profound metabolic changes. Amino acids, such as aspartate, asparagine, serine and alanine, as well as CAC intermediates (e.g. malate, fumarate, succinate) and pyruvate were elevated in 4E-BP1/2 deficient as compared to 4E-BP1/2 proficient cells (Fig. 6A, Fig. S6A). This is consistent with the role of 4E-BP1/2 in suppressing translation of PC, PHGDH, PSAT1, ASNS and TFAM mRNAs (Fig. 5B). The phenformin and lapatinib combination altered levels of the metabolites in both NT2197 control and 4E-BP1/2 KO cells (Fig. 6A, Fig. S6A). However, cells lacking 4E-BP1/2 maintained a significantly higher level of key metabolites, in particular aspartate, asparagine, serine and pyruvate, and to a lesser extent CAC intermediates malate, succinate and fumarate, as compared to control cells even in the presence of the drugs (Fig. 6A, Fig. S6A).

There is mounting evidence that proliferating cells require aspartate and serine synthesis, which are versatile amino acids that are not only involved in protein synthesis, but also other processes including nucleotide synthesis and ROS protection (Lane and Fan, 2015; Vander Heiden and DeBerardinis, 2017). To identify metabolic pathways responsible for the increase in aspartate levels upon ablation of 4E-BP1 and 2, we employed ^13^C-glucose flux analysis. Loss of 4E-BP1/2 increased the proportion out of the total pool of metabolite of m+3 aspartate, malate and fumarate, as compared to control, (Fig. S6B), as well as the level of absolute amount of m+3 aspartate and malate (Fig. 6H, Fig. S6C). This indicates that one mechanism though which cells lacking 4E-BP1/2 maintain high aspartate levels is by using pyruvate to generate malate and aspartate via oxaloacetate (Fig. 6H, Fig. S6B,C). Consistently, cells devoid of 4E-BP1 and 2 expression exhibit elevated levels of pyruvate carboxylase (PC), a rate limiting enzyme that converts pyruvate into oxaloacetate (Fig. 4D), while the lapatinib and biguanide combination has lesser effect on translation of PC mRNA (Fig. 5B).

Addition of 10mM aspartate to the media was sufficient to decrease sensitivity of 4E-BP-proficient cells to lapatinib and phenformin combination to the level observed in 4E-BP1/2-deficient cells (Fig. 6B). In turn, addition of aspartate failed to alter the effects of the drugs in 4E-BP1/2-deficient cells in which aspartate levels are already elevated (Fig. 6B). Taken together, these data suggest that the 4E-BP-dependent regulation of aspartate levels is critical determinant of the efficacy of the phenformin/lapatinib combination.

Aspartate is also essential for *de novo* synthesis of asparagine, and this reaction is catalyzed by asparagine synthetase (ASNS), which simultaneously converts glutamine to glutamate (Balasubramanian et al., 2013). We observed that ASNS mRNA translation and thus ASNS protein levels are downregulated by phenformin and lapatinib combination in 4E-BP1/2-proficient, but not deficient cells (Fig. 5B). Accordingly, cells lacking 4E-BP1/2 contain a higher level of asparagine and glutamate than control cells (Fig. 6A). Moreover, similarly to aspartate, supplementing growth media with asparagine strongly attenuated the anti-proliferative effects of lapatinib and phenformin combination in 4E-BP1/2 proficient cells (Fig. 6C). This suggests that the rescue of proliferation of 4E-BP1/2 proficient cells by aspartate may be mediated by conversion of aspartate into asparagine. Indeed, upon depletion of ASNS, the sensitivity of cells to phenformin and lapatinib was strongly increased (Fig. 6D; Fig. S6D). Accordingly, depletion of asparagine from culture media by L-asparaginase potentiated effects of phenformin/lapatinib combination to a higher extent in 4E-BP-proficient, but not deficient cells (Fig. 6F). Moreover, the importance of ASNS activity in 4E-BP1/2 depleted cells is shown by their increased sensitivity to the phenformin/lapatinib combination following down-regulation of the ASNS protein levels (Fig. 6E, Fig. S6D). Together these findings show that decreased asparagine biosynthesis via inhibition of the mTORC1/4E-BP axis plays a role in synergy between phenformin and lapatinib.

In addition to the effects on aspartate and asparagine synthesis pathway, we observed that enzymes involved in serine biosynthesis pathway PHGDH and PSAT1 are downregulated by the combination of lapatinib and phenformin in 4E-BP1/2 proficient, but not deficient cells (Fig. 4B). We previously demonstrated that serine availability plays a role in determining sensitivity to biguanides (Gravel et al., 2014). Consistently, serine levels and the proportion of m+3 serine derived from ^13^C-glucose are higher in 4E-BP deficient vs proficient cells (Fig. 6A, H; Fig. S6A,C). To further assess the role of serine biogenesis pathway in mediating synergy between lapatinib and phenformin, we depleted PHGDH in 4E-BP1/2 deficient cells. PHGDH depletion increased the anti-proliferative efficacy of lapatinib and phenformin in 4E-BP1/2 deficient cells, therefore indicating that serine biosynthesis pathway also contributes to the anti-proliferative effects of the drug combination (Fig. 6G; Fig. S6E). Collectively these findings demonstrate that the mTORC1/4E-BP axis bolsters proliferation via stimulation of aspartate, asparagine and serine production, and that, though an elevated level of these metabolites, 4E-BP depleted cells are less responsive to the combination between lapatinib and phenformin.

### HIF-1α is a determinant of sensitivity to lapatinib and phenformin

HIF-1α is upregulated in HER2 overexpressing cells and is required for tumor growth (Laughner et al., 2001; Li et al., 2005; Whelan et al., 2013). In NT2197 cells, the phenformin and lapatinib combination treatment strongly decreased HIF-1α levels to an extent that was higher than each drug alone (Fig. 3C). This correlated with a decrease in the levels of mRNAs encoded by HIF-1α target genes (*Vegfa, Glut1, Hk2*; Fig. S7A). mTOR-dependent regulation of HIF-1α levels has been reported to occur by multiple mechanisms including those mediated by 4E-BPs, S6Ks and STAT3 (Dodd et al., 2015). We observed that HIF-1α levels decreased in NT2197 4E-BP1/2-deficient cells treated with lapatinib and phenformin to a comparable degree to 4E-BP1/2-proficient cells (Fig. 4B). Thus, in this context of NT2197 cells, HIF-1α expression is not regulated in a 4E-BP-dependent manner. VHL is a E3 ubiquitin ligase that acts as a principle regulator of the expression of HIF-family members including HIF-1α, leading to their degradation under normoxia (Semenza, 2007). To study the role of HIF-1α in mediating the effects of lapatinib and phenformin, we therefore employed RCC4 renal carcinoma cells which lack VHL and express high levels of HIF-1α protein (Maxwell et al., 1999). RCC4 cells have been reported to be responsive to lapatinib, and lapatinib has been used in RCC clinical trials (Ravaud et al., 2008). The combination of lapatinib and phenformin did not affect HIF-1α levels in VHL-null RCC4 cells (RCC4-mock), as compared to dramatic reduction of HIF-1α in RCC4 cells in which VHL expression was reconstituted (RCC4-VHL) (Fig. 7A). As expected, the hypoxia-mimetic CoCl_2_, which blocks VHL-dependent HIF-1α regulation, abolished the effect of phenformin on HIF-1α in VHL-proficient cells (Fig. S7B). Finally, the effects of the drugs on mTOR signalling were not affected by the VHL status in the cell (Fig. 7A). These findings suggest that the observed reduction in HIF-1α levels induced by lapatinib and phenformin combination are mediated chiefly via VHL, and not via mTOR. RCC4-mock cells were dramatically less sensitive to the phenformin/lapatinib combination relative to RCC4-VHL cells (Fig. 7B). This demonstrates that, in addition to 4E-BPs, VHL-dependent downregulation of HIF-1α critically contributes to biguanide and lapatinib synergy.

**Fig. 7:**
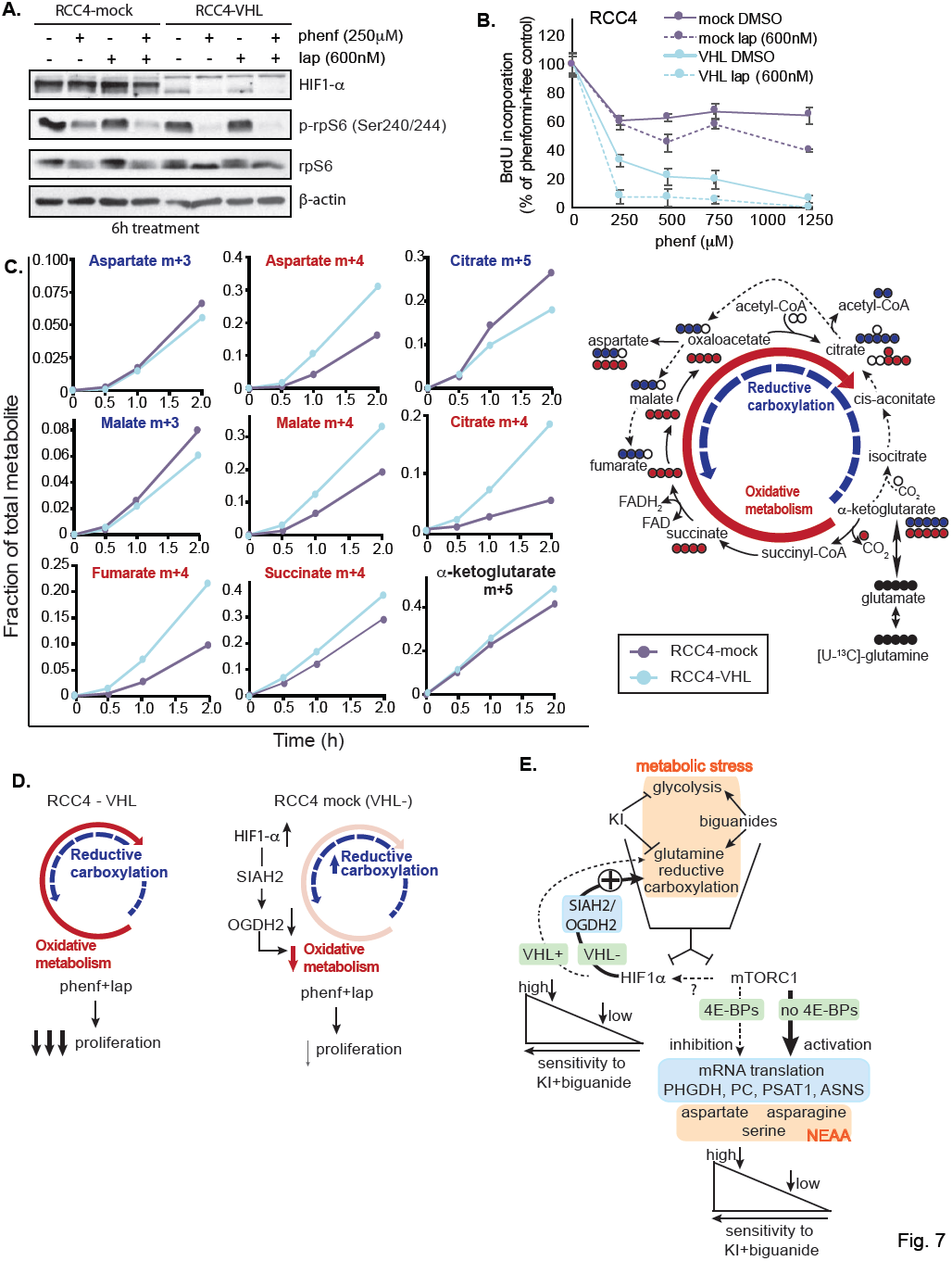
Increase in HIF-1α levels confers partial resistance to phenformin/lapatinib treatment. VHL-null RCC4 cells expressing either empty vector (mock) or VHL-HA were treated with the indicated concentration of phenformin for 6h. Expression of indicated proteins was determined by western blot using appropriate antibodies. β-actin served as a loading control. **B.** VHL-null RCC4 cells expressing either empty vector (mock) or VHL-HA were treated with the indicated concentration of phenformin and lapatinib for 72h. Cell proliferation was measured by BrdU incorporation and normalized to vehicle-treated control. BrdU incorporation values pre-normalization are shown in **Fig. S7C**. The data are presented as mean values +/- the SD (n=3; representative of 3 independent experiments). **C.** VHL-null RCC4 cells expressing either empty vector (mock) or VHL-HA were incubated with ^13^C_5_-glutamine for various time points (30min, 1h, 2h). Stable isotope tracer analyses was performed and the levels of the indicated ion amounts are shown, expressed as fraction of the total pool of the respective metabolite. Data arerepresentative of 2 independent experiments. Bottom right panel: schematic representation of the ^13^C atom incorporation into metabolites. **D**. Schematic representation of the glutamine anapleurotic utilisation in VHL-null RCC4 cells and RCC4-VHL cells. VHL-null RCC4 cells have reduced glutamine oxidative metabolism due to the high HIF1α-, SIAH2-dependent decrease of OGDH2, have increased reductive carboxylation are have reduced sensitivity to phenformin and lapatinib treatment compared to RCC4-VHL cells. **E**. Schematic representation outlining the role of mTORC1/4E-BP axis and HIF1α on the sensitivity of cancer cells to the KI/biguanide combination. The combination of KI and biguanides induces metabolic stress, the inhibition of mTORC1/4E-BP axis and HIF1α reduction in cancer cells, and results in the inhibition of mRNA translation of metabolic genes responsible for the synthesis of key NEAA (aspartate, asparagine, serine). In 4E-BPs depleted cells the increased levels of key NEAA leads to a decreased sensitivity to the drug combination. In VHL-cells, SIAH2-dependent OGDH2 downregulation induces the inhibition of glutamine oxidative metabolism and an increase in glutamine reductive carboxylation, leading to decreased sensitivity to the drug combination.

VHL-null cells have been shown to preferentially engage reductive glutamine metabolism for lipid synthesis (Gameiro et al., 2013; Metallo et al., 2011). Reductive glutamine metabolism appears to be one of the major metabolic adaptations to biguanides (Fendt et al., 2013), and is suppressed when biguanides are combined with lapatinib (Fig. 1D). Indeed, ^13^C-glutamine flux analysis revealed that RCC4-mock cells shunt a larger proportion of glutamine via reductive carboxylation and show a very low activity of glutamine oxidative metabolism compared to RCC4-VHL cells (Fig. 7C). This is indicated by higher levels of citrate (m+5), malate (m+3) and aspartate (m+3) that are generated via reductive glutamine metabolism, in RCC4-mock vs. VHL-RCC4 cells, and *vice versa* higher levels of fumarate (m+4), malate (m+4), aspartate (m+4) and citrate (m+4) which are derived via oxidative glutamine metabolism in RCC4-VHL relative to RCC4-mock cells (Fig. 7C). These findings show that the inhibition of the VHL/HIF-1α pathway and subsequent attenuation of the reductive glutamine mechanism is critical for synergy between biguanides and KIs (Fig. 7D). It has been shown that HIF-1α activity negatively regulates the protein levels of mitochondrial enzyme complex α-ketoglutarate dehydrogenase (αKGDH) subunit E1 (OGDH2) (Sun and Denko, 2014), an enzyme that mediates the oxidation of α ketoglutarate to succinate. Indeed, OGDH2 levels were lower in VHL deficient vs. VHL proficient RCC4 cells (Fig. S7D). This correlated with a decrease in the level of ^13^C-glutamine derived succinate (m+4) in RCC4 mock relative to VHL expressing cells, which is indicative of a reduced capacity for glutamine oxidative metabolism (Fig. 7C,D). Collectively, these data demonstrate that in addition to alterations in NEAA synthesis which are mediated by the mTORC1/4E-BP axis, HIF1α–dependent modulation of reductive glutamine metabolism is also a critical determinant of the efficacy of KI – biguanide combinations (Fig. 7E).

## Discussion

There is heightened interest to identify targetable aspects of the oncometablome. It has been proposed that biguanides may improve efficacy of clinically used KIs. For instance, metformin has been shown to increase the effects of anti-HER2 monoclonal antibody trastuzumab (Herceptin) by suppressing self-renewal and proliferation of cancer stem/progenitor cells in HER2-positive carcinomas (Vazquez-Martin et al., 2011). Similarly, phenformin potentiates the effects of BRAF and ERK inhibitors in BRAF(V600E) and NF1-mutant melanoma cells, respectively, through the selective targeting of JARID1B-expressing subpopulation of cells (Trousil et al., 2017; Yuan et al., 2013). Although that these findings are presently being studied in clinical trials in which biguanides are combined with KIs, the underlying mechanisms and generality of the effects of such combinations are incompletely described. We show that the biguanide phenformin and KIs synergize in distinct cancer cell types that differ not only in their tissue of origin, but also in the driving oncogenic kinase (HER2, EGFR, BRAF or BCR-ABL). Metabolome mapping revealed that the opposing effects of phenformin and KIs on glycolysis are insufficient to explain their synergy. However, perturbations in metabolic and translational landscapes which are mediated via the mTORC1/4E-BP1 axis and HIF-1α are critical for the synergistic effects of KI and biguanide combination.

Our findings reveal mTOR/4E-BP-dependent translational reprograming of mRNAs encoding key enzymes involved in NEAA synthesis including aspartate, asparagine and serine. To this end, inability of the drugs to inhibit the eIF4F assembly results in metabolic adaptations that strongly decrease sensitivity of the cells to KI-biguanide combination (Fig. 7E). Aspartate has been identified as a key metabolite required for the proliferation of cancer cells as it serves as a precursor for both purine and pyrimidine synthesis (Birsoy et al., 2015; Sullivan et al., 2015). Aspartate can be generated from pyruvate though oxaloacetate as via the action of PC, an enzyme mediating the conversion of pyruvate to oxaloacetate (Ling and Keech, 1966). Elevated PC expression and/or activity is observed in breast cancer and non-small-cell lung cancer (NSCLC) (Phannasil et al., 2017; Sellers et al., 2015). We show that PC mRNA translation is regulated via 4E-BPs and that 4E-BP1/2 ablation leads to dramatic increase in aspartate levels and reduction in sensitivity to KI and phenformin combination. Previous reports have underlined the importance of GOT1 for aspartate synthesis in conditions of electron transport chain (ETC) deficiency, an enzyme that mediates the reversible reaction from aspartate to oxaloacetate and from oxaloacetate to aspartate in the cytosol (Birsoy et al., 2015). Phenformin and lapatinib treatment does not alter GOT1 protein levels, nor GOT1 mRNA translation (Fig. 3C, Fig. 5B), suggesting that existence of the alternative mechanism whereby aspartate levels are influenced by enzymatic steps upstream of oxaloacetate synthesis, in particular through the action of PC. Moreover, aspartate conversion to asparagine, which is regulated by 4E-BP-dependent translational regulation of ASNS mRNA, appears to also plays a major role in neoplastic growth and response of malignant cells to phenformin/KI combination. The importance of asparagine for tumor growth is highlighted by the fact that depletion of asparagine pools with L-asparaginase inhibits translation elongation and is used in treating low-ASNS-expressing acute lymphoblastic leukemia and pediatric acute myeloid leukemia (Hill et al., 1967). Moreover, ASNS expression is required for the survival of cultured glioma and neuroblastoma cell lines (Zhang et al., 2014), as well as for the growth of mouse sarcomas (Hettmer et al., 2015). Asparagine levels were also shown to function in the regulation of amino-acid uptake, and to positively regulate the expression of genes of the serine synthesis pathway (Krall et al., 2016).

In addition to aspartate synthesis, we show that mTORC1/4E-BP axis plays a major role in serine biogenesis by modulating translation of PHGDH and PSAT1 mRNAs. Accordingly, serine synthesis from ^13^C labeled glucose is increased in 4E-BP depleted cells compared to control, whilst PHGDH knock-down increased the sensitivity to phenformin and KI treatment in both control and 4E-BP depleted cells. Serine biosynthesis has been documented to be essential for neoplastic growth (Mattaini et al., 2016). Cancer cells exhibit increased serine biosynthesis (Davis et al., 1970; Snell, 1984), PHGDH and PSAT1 are overexpressed in breast cancer, melanoma, cervical cancer, glioma (Locasale et al., 2011; Possemato et al., 2011) and characterize poor-prognosis for non–small cell lung cancers and lung adenocarcinoma ((DeNicola et al., 2015; Zhang et al., 2017). Increase in serine synthesis pathway flux appears to be beneficial for cancer cells irrespective of the availability of serine in the environment, or intracellular serine levels (Mattaini et al., 2016). This is explained by the tenet that serine is a central node for the biosynthesis of many metabolites (e.g. nucleotides, folate metabolism) (Lehninger et al., 2000; Locasale, 2013), that it can facilitate amino acid transport, as well as redox homeostasis (Fan et al., 2014) (Mattaini et al., 2016). Accordingly, herein we show that 4E-BP-dependent translational regulation of serine synthesis is a critical determinant of sensitivity of cancer cells to biguanides and kinase inhibitor combinations.

In addition to 4E-BPs-dependent metabolic reprograming, we show that HIF-1α-dependent modulation of reductive glutamine metabolism plays a central role in synergy between KIs and phenformin. Cancer cells driven by HER2 or BCR-ABL upregulate HIF-1α, even under normoxic conditions (Laughner et al., 2001; Li et al., 2005; Mayerhofer et al., 2002; Semenza, 2013). Increased HIF-1α expression is required for neoplastic growth of HER2+ cancer cells (Whelan et al., 2013), survival of chronic myeloid leukemia stem cells (Zhang et al., 2012), and is associated with the metabolic reprograming in BCR-ABL inhibitor resistant cells (Zhao et al., 2010). The decrease in HIF-1α levels induced by KI - biguanide combinations appears to occur chiefly via VHL, which is in accordance with previous reports showing that metformin induces HIF-1α degradation through inhibition of mitochondrial complex I (Wheaton et al., 2014). RCC4 cells which lack VHL and thus exhibit high HIF-1α levels showed decreased sensitivity to KI-biguanide combination, which coincided with high rates of reductive glutamine metabolism and decrease in glutamine oxidative metabolism. This reduction in glutamine oxidative metabolism in VHL deficient, high HIF1αcells is mediated by the SIAH2-dependent degradation of αketoglutarate dehydrogenase subunit OGDH2 (Sun and Denko, 2014). Notably, although phenformin and lapatinib combination reduced HIF1α protein levels in VHL proficient cells, it also decreased OGDH2 levels. This is however consistent with the inhibition of OGDH by NADH (Strumilo, 2005) in the context of ETC deficiency, as the inhibition of mitochondrial complex I (NADH dehydrogenase) by biguanides is accompanied by an increased accumulation of NADH. These data show that a second genetic modification, VHL loss, and the resulting activation of HIF-1α allows cells to overcome the antineoplastic effects of the biguanide-KI combination (Fig. 7E). Therefore, the mTORC1/4E-BP and VHL/HIF-1α axes constitute two independent mechanisms which allow cells to adapt to KI-biguanide combinations.

Strong mTOR inhibition by active site mTOR inhibitors is associated with metabolic dormancy, due to concomitant reductions in both mitochondrial ATP production and cellular ATP consumption by translational machinery and other anabolic processes (Gandin et al., 2016; Morita et al., 2013; Morita et al., 2015). In stark contrast, KI-phenformin combinations, although inducing considerably less inhibition of mTOR and 4E-BP phosphorylation, result in cell death. Our findings therefore suggest that mild mTOR inhibition which is sufficient to induce translational reprograming of the metabolome, but not to decrease energy consumption is paradoxically more likely to lead to an energetic crisis and cell death than complete mTOR inhibition. This suggests that targeting oncogenic mTOR signaling with combinations of upstream KIs and biguanides may be an alternative to the current approaches in the clinic employing mTOR inhibitors. It is however important to note that early results of randomized, placebo-controlled clinical trials of biguanides in oncology have been disappointing (Kordes et al., 2015). One reason for this may relate to sub-optimal pharmacokinetics of metformin (Chandel et al., 2016; Dowling et al., 2016), an issue which may be addressed by the use of novel biguanides under development (Choi et al., 2016; Zhang et al., 2016). Other trials, including one using phenformin in combination with BRAF inhibitors are underway (ClinicalTrials.gov; NCT03026517).

In conclusion, we show that 4E-BP-dependent translational regulation of biosynthesis of NEAA serine, aspartate and asparagine, in combination with HIF-1α dependent switch to reductive glutamine metabolism plays a major role in energy stress response of cancer cells and underlies the sensitivity to KI and biguanide combinations. Importantly, these findings show that drug-induced energy stress results in rapid adaptation of the oncometabolome. This plasticity of the oncometabolome represents a paramount challenge, as genetic and/or epigenetic alterations, including those affecting 4E-BP and HIF-1α function, enable cancer cells to evade metabolic stresses and may thus hinder efficacy of the future metabolic cancer therapies.

## Materials and Methods

### Cell culture and inhibitors

NMuMG-NT2197 cells were cultured in Dulbecco’s modified Eagle’s medium (DMEM; Wisent #319-005-CL), supplemented with 10% heat-inactivated fetal bovine serum (Wisent), 1% penicillin/streptomycin (Wisent; #450-201-EL) and 2mM L-Glutamine (Wisent, #609-065-EL) to obtain 6mM L-Glutamine final concentration, 10 μ g/ml insulin, 20 mM HEPES, pH 7.5 (Wisent, #330-050-EL), and were a generous gift from Dr. Ursini-Siegel (McGill University) (Ursini-Siegel et al., 2008). A375 and HCT116 cells were maintained in DMEM supplemented with 10% fetal bovine serum (Wisent), 1% penicillin/streptomycin (Wisent; #450-201-EL) and 2mM L-Glutamine (Wisent, #609-065-EL) to obtain 6mM L-Glutamine final concentration, and were a generous gift from Dr. Ronai (*Sanford-Burnham* Medical Research Institute) and Dr. Zhdanov (Univ. College Cork), respectively. K562 cells (ATTC) were cultured in RPMI-1640 (Wisent), supplemented with 10% fetal bovine serum (Wisent) and 1% penicillin/streptomycin (Wisent; #450-201-EL). RCC4 mock or expressing VHL-HA were cultured in DMEM supplemented with 10% fetal bovine serum (Wisent), 1% penicillin/streptomycin (Wisent; #450-201-EL) and 250μM G418. Where indicated, cells were treated with lapatinib ditosylate (Selleckchem), plx4032 (Selleckchem), imatinib mesylate (Selleckchem), phenformin hydrochloride (Sigma), Torin1 (Tocris Bioscience), 3-PO (Calbiochem), L-aspartic acid (Sigma), L-asparagine (Sigma), or asparaginase (SMBD JGH pharmacy; 5000units/ml stock).

All lentiviral shRNA vectors were retrieved from the arrayed Mission TRC genome-wide shRNA collections purchased from Sigma-Aldrich Corporation. Additional information about the shRNA vectors can be found at http://www.sigmaaldrich.com/life-science/functional-genomics-andrnai/hrna/library-information.html or http://www.broad.mit.edu/genome_bio/trc/rnai.html, using the TRCN number. The following lentiviral shRNA vector were used: TRCN0000041627 (mPHGDH), TRCN0000324779 (mASNS). The Non-Target shRNA Control (Sigma: SHC002) was used as negative control. Lentiviral supernatants were generated as described at http://www.broadinstitute.org/rnai/public/resources/protocols. Supernatants were applied on target cells with polybrene (6 μg/mL). Cells were reinfected the next day and, 2 days later, selected with puromycin for 72 hours (4 μg/mL; Sigma).

### Proliferation and cell viability assays

For the bromodeoxyuridine (BrdU) incorporation assay (Cell Proliferation ELISA BrdU Kit from Roche), cells were seeded in 96-well plates (1000 cells/well) and maintained as indicated in the figure legends of Fig. 1, Fig. 2, Fig. 4, Fig. 6, Fig. 7, and Fig. S1, Fig. S2, Fig. S4 and Fig. S7 for 72h. Tests were performed in triplicate, with each microplate including media and DMSO control wells, as per the manufacturer’s instruction. Absorbance at 370nm (reference wavelength 492nm) was measured using a Benchmark Plus microplate reader (Bio-Rad). Combination index_50_ (CI_50_) was calculated using the isobologram equation (CI_50_) = (*a/A*) + (*b/B*), where, *A* and *B* represent the IC_50_ concentrations of KI (lapatinib, PLX4232 or imatinib) and phenformin when administered alone, *a* and *b* represent the concentrations required for the same effect when KI and phenformin were administered in combination, and CI_50_ represents an index of drug interaction (combination index). CI_50_ values of < 1 indicate synergy, a CI_50_ value of 1 represents additivity, and CI_50_ values of > 1 indicate antagonism (Berenbaum, 1989; Chou and Talalay, 1984; Tallarida, 2006). Data points shown were calculated from 2 independent experiments.

### Xenograft experiment

NMuMG/NT2197 mammary tumor cells (5×10^4^) were injected into both sides of the fourth mammary fat pads of Nude mice (Charles River Laboratories). The mammary tumor growth (mm^3^ ± SEM) was obtained by caliper measurements (n=10 tumors/cell line) and were calculated according to following formulae: 4/3π(w^2^ × l) as previously described, where l and ware diameter measurements of the length and width of the tumor (Ursini-Siegel et al., 2007). Once tumor volume reached 100∼200 mm^3^, mice were given vehicle control, phenformin (50mg/kg) (Sigma-Aldrich P7045) via intraperitoneal injection, and/or lapatinib (50mg/kg) (lapatinib: LC Laboratories L-4804) via gavage every 24hr. All treatment conditions contained 12.5% DMSO and were resuspended in 0.5% methylcellulose-0.1% Tween-80 solution (Tween-80: Sigma-Aldrich P4780, methyl cellulose: 274429). On the day of necropsy mice were sacrificed 4h post-treatment, the mammary tumors were placed in 10% formalin to be fixed and later embedded in paraffin for immunohistochemistry.

### Immunohistochemistry

Immunohistochemical staining of paraffin-embedded sections was performed as described previously (Ursini-Siegel et al., 2008), using the following antibodies: phospho-rpS6 (1:2000, CST 2215), phopsho-AMPK (1:100, CST 2535), Ki67 (1:500, Abcam ab15580) and cleaved Caspase3 (1:250, CST 9661). Quantification of stained sections was performed using Aperio Imagescope software.

### Western blotting and antibodies

Cells were scraped in ice-cold PBS (pH 7.4), washed by centrifugation (800xg/5 minutes at 4°C) and lysed on ice in [50mM Tris/HCL pH 7.4, 5mM NaF, 5mM Na pyrophosphate, 1mM EDTA, 1mM EGTA, 250mM mannitol, 1% (v/v) triton x-100, 1mM DTT, 1X complete protease inhibitors (Roche), 1X PhosSTOP (Sigma-Aldrich)] for 30 minutes with occasional vortexing. After lysis, cellular debris was removed by centrifugation at (16,100xg/20 minutes at 4°C). Protein concentrations in cell extracts were determined using Pierce BCA Protein Assay Kit (Thermo Fisher Scientific). Protein extracts (10-40 μ g of protein) were resuspended in Laemmli buffer, heated to 95°C and separated by sodium dodecyl sulfate-polyacrylamide gel electrophoresis and transferred to nitro-cellulose membrane (Bio-Rad). To detect proteins of interest we used the following primary antibodies: anti-β-actin (Clone AC-15) #A1978, anti-HA #H6908, anti-PHGDH #HPA021241, from Sigma (Saint Louis, Missouri, USA); anti-4E-BP1(53H11) #9644, anti-p-4E-BP1 (Ser65) (174A9) #9456, anti-4E-BP2 #2845, anti-p-rpS6 (Ser240/244) #2215, anti-S6K #2708, anti p-S6K (Thr389) (#9234), anti ACC #3662, anti p-ACC (Ser79) #3661, anti ERK1/2 #9102, anti p-ERK1/2 (Thr202/Tyr204) #9101, anti eIF2α (L57A5) #2103, anti p-eIF2 α (Ser51) #9721, anti c-myc #9402, anti-Cyclin D3 (DCS22) #2936, anti-eIF4G1 #2858, anti-BCL-2 #2870, anti-TFAM #7495, anti-GAPDH (14C10) #2118, all from Cell Signaling Technologies (Danvers, MA, USA); anti-rpS6 (C-8) #sc-74459, anti-MCL-1(H-260) #sc-20679, anti-pyruvate carboxylase (PCB; H-300) #67021, from Santa Cruz Biotechnologies (Dallas, Texas, USA); anti-eIF4E #610269 from BD Biosciences (San Jose, CA, USA); anti-survivin #SURV11-S from Alpha Diagnostics (San Antonio, Texas, USA); anti-ASNS #NBP1-87444, anti-PSAT1 #NBP1-32920, from Novus Biologicals; HIF1α # 10006421 from Cayman Chemical; and from anti-OGDH #ab87057, anti-GOT1 #ab189863, anti-survivin #ab175809, anti-TFAM #ab131607, all from Abcam. All primary antibodies were incubated in 5% (w/v) BSA in 1XTBS/0.5%Tween20 (Sigma) over night at 4°C. Secondary anti-mouse-HRP and anti-rabbit-HRP antibodies (Amersham) were used at 1:5,000 dilution in 5% (w/v) non-fat dry milk in 1XTBS/0.5%Tween 20 for 1 hour at room temperature. Signals were revealed by chemiluminescence (ECL, GE Healthcare) on HyBlot CL autoradiography film (Denville Scientific Inc. #E3018).

### In vitro cap-binding affinity assay

Experiments were carried out as previously described (Dowling et al., 2010). Briefly, cells were seeded in 150mm plates and treated as indicated in Fig. 4 for 4h. Cells were washed with cold PBS, collected, and lysed in buffer containing 50mM MOPS/KOH (7.4), 100mM NaCl, 50 mM NaF, 2mM EDTA, 2mM EGTA, 1% NP40, 1% sodium deoxycholate, 7 mM β-mercaptoethanol, 1X complete protease inhibitors (Roche), and 1X PhosSTOP (Sigma-Aldrich). Lysates were incubated with m^7^-GDP-agarose beads (Jena Bioscience) for 20 min, washed 4 times with the washing buffer containing 50mM MOPS/KOH (7.4), 100mM NaCl, 50 mM NaF, 0.5mM EDTA, 0.5 mM EGTA, 7 mM β-mercaptoethanol, 0.5 mM PMSF, 1mM Na_3_VO_4_ and 0.1mM GTP, and bound proteins were eluted by boiling the beads in loading buffer. eIF4F complex formation was assessed by western blot of eluted proteins, using 4E-BP1, eIF4G1, eIF4E and β-actin antibodies as described in the “Western blot analysis and antibodies” section.

### Proximity ligation assay (PLA)

Interactions between eIF4E and eIF4G1 (eIF4E–eIF4G) or eIF4E and 4E-BP1 (eIF4E–4E-BP1) were detected by *in situ* proximity ligation assay (PLA) on NMuMG-NT2197 cells according to the manufacturer’s instructions (Duolink In situ red mouse/rabbit kit, Sigma). Briefly, 1.5×10^ 5 NT2197 control or 4E-BP1/2 DKO cells were seeded in six-well plates containing coverslips. The day after, cells were treated for 4 hours either with DMSO or combination of lapatinib (600nM) and phenformin (250μ M), washed with PBS and fixed in 3.7% formaldehyde for 15 min at room temperature. Cells were permeabilized for 10 min in PBS containing 0.2% Triton X-100, blocked for 45 min in PBS containing 10% FBS (blocking buffer) and incubated with primary antibodies (anti-eIF4E from Santa Cruz Biotechnologies SC-271480, 1:200; anti-eIF4G1 from Cell Signaling Technology #2498, 1:200 and anti-4E-BP1 from Cell Signaling Technology #9644, 1:500) overnight at 4°C. Secondary antibodies conjugated with PLA minus and PLA plus probes were incubated for 1 hour at 37°C. Ligation was performed for 30min at 37°C followed by amplification with polymerase for 2 hours at 37°C. Wheat germ agglutinin conjugated to Alexa fluor-488 (WGA, Life technologies), used to stain the plasma membrane, was incubated for 20 min at room temperature. Coverslips were mounted with O-link mounting medium containing DAPI (Sigma). Immunofluorescence microscopy images from single confocal sections were acquired at room temperature with a Wave FX spinning disc confocal microscope (Leica) using a 40X oil-immersion objective. All images were acquired using identical parameters. 10 images per condition were analyzed. Foci corresponding to eIF4E-eIF4G1 or eIF4E-4E-BP1 interactions and nuclei were detected and quantified using the Log-Dog Spot Counter in ImageJ (BAR, Ferreira et al (2016). DOI: 10.5281/zenodo.28838). The number of PLA foci per cell were log2-transformed and represented as fold-change to DMSO. P-values were calculated using one-way ANOVA (for eIF4E/4E-BP1) and Tukey test (for eIF4E–eIF4G1).

### Flow cytometry analysis of apoptotic cells

NMuMG-NT2197 cells were plated in 6-well plates and grown for 24 h, after which they were treated, as indicated in Fig. 1 and Fig. 4, with DMSO control, lapatinib, phenformin or both for 72h. Cells were trypsinized, counted and 100,000 cells were stained using Annexin V-FITC and PI for 20min in the dark as per the manufacturer’s instructions (FITC Annexin V Apoptosis Detection Kit, BD Biosciences). Samples were analyzed with a LSRII cytometer (Becton-Dickinson, Mountain View, CA). Fluorescence was detected by excitation at 488nm and acquisition on the 530/30-A channel for FITC-Annexin V and by excitation at 561nm and acquisition on the 610/20-A channel for PI. Cell populations were separated as follows: viable cells – Annexin V-/PI-; early apoptosis - Annexin V+/PI-; late apoptosis Annexin V+/PI+ and expressed as % of total single cells. Experiments were carried out 3 times independently (n=3), with 2 technical replicates in each experiment.

### Flow cytometry analysis of cytotoxicity

K562 cells were seeded in 6-well plates and grown for 24 h, after which they were treated, as indicated in Fig. S1, with DMSO control, lapatinib, phenformin or both for 72h. Cells were counted and 100,000 cells were stained using DAPI (4′,6-diamidino-2-phenylindole; 0.5 μ g/ml) for 15min in the dark. Samples were analyzed with a LSRII cytometer (Becton-Dickinson, Mountain View, CA). Fluorescence was detected by excitation at 405nm and acquisition on the 450/50-A channel. Cell populations with medium and high intensity DAPI stain were quantified. Experiments were carried out 3 times independently (n=3), with 2 technical replicates in each experiment.

### Generation of 4E-BP1/2 knock-out cells by CRISR-Cas9

A375 cells were transfected in 6 well plates with a plasmid expressing hCas9 (0.7μg); three guides RNAs (gRNAs) targeting h4E-BP1 (purchased from GeneCopoeia, cat. # HCP204676-SG01-3-B-a; HCP204676-SG01-3-B-b; and HCP204676-SG01-3-B-c against h4E-BP1 sequences CCGCCCGCCCGCTTATCTTC; GTGAGTTCCGACACTCCATC; and TGAAGAGTCACAGTTTGAGA, respectively); three gRNAs targeting h4E-BP2 (purchased from GeneCopoeia, cat. # HCP254214-SG01-3-B-a; HCP254214-SG01-3-B-b; and HCP254214-SG01-3-B-c against h4E-BP2 sequences GTGGCCGCTGCCGGCTGACG; CTAGTGACTCCTGGGATATT; and ACAACTTGAACAATCACGAC); (0.2μg for each gRNA to total 1.2μg); and pBabe-puro (0.6μg), using Lipofectamine 2000 (Invitrogen), according to the manufacturers instruction. As a control, cells were transfected with a plasmid expressing hCas9 (0.7μg) and pBabe-puro (0.6μg), using Lipofectamine 2000 (Invitrogen). NT2197 cells were transfected in 6 well plates with a plasmid expressing hCas9 (0.7μg); three gRNAs targeting m4E-BP1 (purchased from GeneCopoeia, cat. # MCP227000-SG01-3-B-a; MCP227000-SG01-3-B-b; and MCP227000-SG01-3-B-c against m4E-BP1 sequences GAGCTGCACGCCATCGCCGA;CGTGCAGGAGACATGTCGGC; an dGACTACAGCACCACTCCGGG, respectively); three gRNAs targeting m4E-BP2 (purchased from GeneCopoeia, cat. # MCP229088-SG01-3-B-a; MCP229088-SG01-3-B-b; and MCP229088-SG01-3-B-c against m4E-BP2 sequences GGAGCCATGTCCGCGTCGGC; CTGATAGCCACGGTGCGCGT; and GCGCCATGGGAGAATTGCGA); (0.2μg for each gRNA to total 1.2μg); and pQCXIB(blast) (0.6μg), using Lipofectamine 2000 (Invitrogen), according to the manufacturers instruction. As a control, cells were transfected with a plasmid expressing hCas9 (0.7μg) and pQCXIB(blast) (0.6μg), using Lipofectamine 2000 (Invitrogen). Two days post-transfection, A375 cells were selected for 3 days with puromycin (4μg/ml) and NT2197 were selected for 3 days with blasticidin (8μg/ml), to remove non-transfected cells. Following selection, NT2197 and A375 cells were seeded in 96 well plates at a density of single cell/well in puromycin or blasticidin free media. Cells were monitored to the presence of single colonies/well. Single cell colonies were amplified to generate cell lines and the expression of 4E-BP1 and 4E-BP2 was analysed by western blot. Lines with loss of 4E-BP1 and 4E-BP2 expression were kept for further experiments. For the control cells, single cell colonies were amplified and 5 of the control lines were pooled to generate the A375 or NT2197 CRISPR control population.

### Polysome profiles and RT-qPCR

Briefly, NT2197 CTR and 4E-BP1/2 CRISPR depleted cells were seeded in two 15cm Petri dish, harvested at 80% confluency and lysed in hypotonic lysis buffer (5mM Tris HCl pH7.5, 2.5mM MgCl_2_, 1.5mM KCl, 100 μg/ml cycloheximide, 2mM DTT, 0.5% Triton, 0.5% Sodium Deoxycholate). Polysome-profiling was carried out as described (Gandin et al., 2014). Fractions were collected as described in (Gandin et al., 2014) and RNA was extracted using Trizol (Thermo Fisher Scientific) according to manufacturer’s instructions. RT-qPCR was performed as previously described (Miloslavski et al., 2014). RT and qPCR were performed by SuperScript III Reverse Transcriptase and Fast SYBR Green Mastermix (Invitrogen), respectively. Experiments were done at least in independent duplicates (n=2) whereby every sample was analyzed in a technical triplicate. Analyses were carried out using relative standard curve method as described in http://www3.appliedbiosystems.com/cms/groups/mcb_support/documents/generaldocuments/cms_040980.pdf. Primers are listed in the **Supplemental item 3**. Polysome to 80S ratios were calculated by comparing the area under the 80S peak and the combined area under the polysome peaks.

### Glucose consumption assays

NT2197 and A375 cells were seeded in 6-well plates and cultured in complete medium for 24h in order to obtain 30% density. Medium was replaced with 2ml of indicated treatment media, and cells were cultured for 22h. Treatment media was replaced with 1 ml of fresh treatment media for an additional 4h. Subsequently, supernatant samples were collected and cells were counted using an automated cell counter (Invitrogen). For K562, cells were seeded in 6-well plates and cultured in treatment media for 24h. Subsequently, supernatant samples were collected and cells were counted using an automated cell counter (Invitrogen). Measurement of glucose concentration in samples was done as previously described (Blake and McLean, 1989). Total consumption was calculated by subtracting results from baseline glucose concentration, measured in samples from media incubated in identical conditions, without cells. Molar concentrations of glucose were multiplied by total media volume/well (1ml) and normalized per 10^ 6 cells; data were expressed as μM glucose/10^ 6cells/h.

### GC/MS and stable isotope tracer analyses

NT2197, A375 and RCC4 cells were cultured in 6-well plates (9.6 cm^2^/well) to 80% confluency, after which indicated treatment was added to the media for 18 hours. For stable isotope tracer analyses, the media was then exchanged for 10mM [U-^13^C]-glucose (CLM-1396-PK; Sigma)-labeled media or 4mM [U-^13^C]-glutamine (CLM-1822-H-PK; Sigma)-labeled media for various periods from 3min to 2h, as indicated. Cells were then rinsed three times with 4°C saline solution (9 g/L NaCl) and quenched with 600 μl 80% MEOH (< 20°C). For total level of metabolites (non-labeled, steady-state), cells were rinsed three times with 4°C saline solution (9 g/L NaCl) and quenched with 300 μl 80% methanol (< 20°C). Cells were removed from plates and transferred to prechilled tubes. MeOH quenching and harvest were repeated for a second time to ensure complete recovery. The remaining procedure is identical for steady state and stable isotope tracer analysis. Cell lysis was carried by sonication at 4°C (10 minutes, 30 sec on, 30 sec off, high setting, Diagenode Bioruptor). Extracts were cleared by centrifugation (14,000 rpm, 4°C) and supernatants were dried in a cold trap (Labconco) overnight at −4°C. Pellets were solubilized in 30 μl pyridine containing methoxyamine-HCl (10 mg/ mL) by sonication and vortex, centrifuged and pellets were discarded. Samples were incubated for 30 minutes at 70°C (methoximation), and then were derivatized with MTBSTFA (70 μl, Sigma) at 70°C for 1 h. One μL was injected per sample for GC–MS analysis. GC–MS instrumentation and software were all from Agilent. GC– MS methods and mass isotopomer distribution analyses are as previously described (Gravel et al., 2016; Morita et al., 2013). Data analyses were performed using the Chemstation software (Agilent, Santa Clara, USA).

### Statistical analyses

Data shown for BrdU incorporation, apoptosis, glucose uptake, and PLA are derived from n=3 technical replicates and are representative of three independent experiments. Data shown for western blotting are representative of three independent experiments. Data shown for polysome profiling/qPCR and metabolic experiments are derived from n=3 technical replicates and are representative of two independent experiments.

Statistical analysis was performed using two-way ANOVA (unless otherwise specified). For Fig. 6F, an analysis of variance using R software was fit to test for interaction between cell line and the asparaginase dose, and including also a factor for experiment. The test for interaction was significant (p=0.00803). Predictions and confidence intervals, taken from the model (with experiment=1) showed that the confidence intervals for conditions B (CTR asparaginase, phenformin and lapatinib) and D (4E-BP DKO cl17 asparaginase, phenformin and lapatinib) did not overlap and that "OD.blank" was substantially lower in condition B when compared to D, although both " OD.blank" values were extremely similar when the asparaginase dose was zero (i.e. comparing A and C, CTR no asparaginase and 4E-BP DKO cl17 no asparaginase, respectively).

## Acknowledgements

The authors thank Yunhua Zhao for technical support, and Shawn McGuirk for the conceptual design of tracing schematics. IT is supported by Junior 2 FRQ-S award. JUS is supported by a Senior FRQ-S award. MC is recipient of the CIHR Postdoctoral fellowship. This research was funded by grants Canadian Cancer Society Research Institute (CCSRI-703816) to IT and MP, Canadian Institutes for Health Research (MOP-363027) to IT and (MOP-111143) to JUS, Terry Fox Research Institute (TFF-116128) to IT, MP and J.S-P. The Rosalind and Morris Goodman Cancer Research Centre Metabolomics Core Facility is supported by the Canada Foundation for Innovation, The Dr. John R. and Clara M. Fraser Memorial Trust, the Terry Fox Foundation (TFF Oncometabolism Team Grant 1048 in partnership with the Foundation du Cancer du Sein du Quebec), and McGill University.

## References

Algire, C., Amrein, L., Bazile, M., David, S., Zakikhani, M., and Pollak, M. (2011). Diet and tumor LKB1 expression interact to determine sensitivity to anti-neoplastic effects of metformin in vivo. Oncogene 30, 1174–1182.

Andrzejewski, S., Gravel, S. P., Pollak, M., and St-Pierre, J. (2014). Metformin directly acts on mitochondria to alter cellular bioenergetics. Cancer Metab 2, 12.

Anisimov, V. N., Berstein, L. M., Egormin, P. A., Piskunova, T. S., Popovich, I. G., Zabezhinski, M. A., Kovalenko, I. G., Poroshina, T. E., Semenchenko, A. V., Provinciali, M., et al. (2005). Effect of metformin on life span and on the development of spontaneous mammary tumors in HER-2/neu transgenic mice.Exp Gerontol 40, 685–693.

Assan, R., Heuclin, C., Girard, J. R., LeMaire, F., and Attali, J. R. (1975). Phenformin-induced lactic acidosis in diabetic patients. Diabetes 24, 791–800.

Bailey, C. J., and Turner, R. C. (1996). Metformin. N Engl J Med 334, 574–579.

Balasubramanian, M. N., Butterworth, E. A., and Kilberg, M. S. (2013). Asparagine synthetase: regulation by cell stress and involvement in tumor biology. Am J Physiol Endocrinol Metab 304, E789–799.

Beloueche-Babari, M., Box, C., Arunan, V., Parkes, H. G., Valenti, M., De Haven Brandon, A., Jackson, L. E., Eccles, S. A., and Leach, M. O. (2015). Acquired resistance to EGFR tyrosine kinase inhibitors alters the metabolism of human head and neck squamous carcinoma cells and xenograft tumours. Br J Cancer 112, 1206–1214.

Ben Sahra, I., Laurent, K., Giuliano, S., Larbret, F., Ponzio, G., Gounon, P., Le Marchand-Brustel, Y., Giorgetti-Peraldi, S., Cormont, M., Bertolotto, C., et al. (2010). Targeting cancer cell metabolism: the combination of metformin and 2-deoxyglucose induces p53-dependent apoptosis in prostate cancer cells. Cancer Res 70, 2465–2475.

Ben Sahra, I., Laurent, K., Loubat, A., Giorgetti-Peraldi, S., Colosetti, P., Auberger, P., Tanti, J. F., Le Marchand-Brustel, Y., and Bost, F. (2008). The antidiabetic drug metformin exerts an antitumoral effect in vitro and in vivo through a decrease of cyclin D1 level. Oncogene 27, 3576– 3586.

Ben Sahra, I., Regazzetti, C., Robert, G., Laurent, K., Le Marchand-Brustel, Y., Auberger, P., Tanti, J. F., Giorgetti-Peraldi, S., and Bost, F. (2011). Metformin, independent of AMPK, induces mTOR inhibition and cell-cycle arrest through REDD1. Cancer Res 71, 4366–4372.

Berenbaum, M. C. (1989). What is synergy? Pharmacological reviews 41, 93–141.

Bhat, M., Robichaud, N., Hulea, L., Sonenberg, N., Pelletier, J., and Topisirovic, I. (2015). Targeting the translation machinery in cancer. Nat Rev Drug Discov 14, 261–278.

Birsoy, K., Wang, T., Chen, W. W., Freinkman, E., Abu-Remaileh, M., and Sabatini, D. M. (2015). An Essential Role of the Mitochondrial Electron Transport Chain in Cell Proliferation Is to Enable Aspartate Synthesis. Cell 162, 540–551.

Blake, D. A., and McLean, N. V. (1989). A colorimetric assay for the measurement of D-glucose consumption by cultured cells. Anal Biochem 177, 156–160.

Brattain, M. G., Fine, W. D., Khaled, F. M., Thompson, J., and Brattain, D. E. (1981). Heterogeneity of malignant cells from a human colonic carcinoma. Cancer Res 41, 1751–1756.

Bridges, H. R., Jones, A. J., Pollak, M. N., and Hirst, J. (2014). Effects of metformin and other biguanides on oxidative phosphorylation in mitochondria. Biochem J 462, 475–487.

Brodaczewska, K. K., Szczylik, C., Fiedorowicz, M., Porta, C., and Czarnecka, A. M. (2016). Choosing the right cell line for renal cell cancer research. Mol Cancer 15, 83.

Brunn, G. J., Hudson, C. C., Sekulic, A., Williams, J. M., Hosoi, H., Houghton, P. J., Lawrence, J. C., Jr., and Abraham, R. T. (1997). Phosphorylation of the translational repressor PHAS-I by the mammalian target of rapamycin. Science 277, 99–101.

Buttgereit, F., and Brand, M. D. (1995). A hierarchy of ATP-consuming processes in mammalian cells. Biochem J 312 (Pt 1), 163-167.

Cerezo, M., Tichet, M., Abbe, P., Ohanna, M., Lehraiki, A., Rouaud, F., Allegra, M., Giacchero, D., Bahadoran, P., Bertolotto, C., et al. (2013). Metformin blocks melanoma invasion and metastasis development in AMPK/p53-dependent manner. Mol Cancer Ther 12, 1605–1615.

Chandel, N. S., Avizonis, D., Reczek, C. R., Weinberg, S. E., Menz, S., Neuhaus, R., Christian, S., Haegebarth, A., Algire, C., and Pollak, M. (2016). Are Metformin Doses Used in Murine Cancer Models Clinically Relevant? Cell Metab 23, 569–570.

Choi, J., Lee, J. H., Koh, I., Shim, J. K., Park, J., Jeon, J. Y., Yun, M., Kim, S. H., Yook, J. I.,Kim, E. H., et al. (2016). Inhibiting stemness and invasive properties of glioblastoma tumorsphere by combined treatment with temozolomide and a newly designed biguanide (HL156A). Oncotarget 7, 65643–65659.

Chou, T. C., and Talalay, P. (1984). Quantitative analysis of dose-effect relationships: the combined effects of multiple drugs or enzyme inhibitors. Advances in enzyme regulation 22, 27-55.

Clem, B., Telang, S., Clem, A., Yalcin, A., Meier, J., Simmons, A., Rasku, M. A., Arumugam, S., Dean, W. L., Eaton, J., et al. (2008). Small-molecule inhibition of 6-phosphofructo-2-kinase activity suppresses glycolytic flux and tumor growth. Mol Cancer Ther 7, 110–120.

Davis, J. L., Fallon, H. J., and Morris, H. P. (1970). Two enzymes of serine metabolism in rat liver and hepatomas. Cancer Res 30, 2917–2920.

De Benedetti, A., Joshi, B., Graff, J. R., and Zimmer, S. G. (1994). CHO cells transformed by the translation factor elF-4E display increased c-Myc expression, but require overexpression of Max for tumorigenicity. Mol cell Diff 2, 347–371.

DeBerardinis, R. J., and Chandel, N. S. (2016). Fundamentals of cancer metabolism. Sci Adv 2, e1600200.

Deblois, G., Smith, H. W., Tam, I. S., Gravel, S. P., Caron, M., Savage, P., Labbe, D. P., Begin, L. R., Tremblay, M. L., Park, M., et al. (2016). ERRalpha mediates metabolic adaptations driving lapatinib resistance in breast cancer. Nat Commun 7, 12156.

DeNicola, G. M., Chen, P. H., Mullarky, E., Sudderth, J. A., Hu, Z., Wu, D., Tang, H., Xie, Y., Asara, J. M., Huffman, K. E., et al. (2015). NRF2 regulates serine biosynthesis in non-small cell lung cancer. Nature genetics 47, 1475–1481.

Dirat, B., Ader, I., Golzio, M., Massa, F., Mettouchi, A., Laurent, K., Larbret, F., Malavaud, B., Cormont, M., Lemichez, E., et al. (2015). Inhibition of the GTPase Rac1 mediates the antimigratory effects of metformin in prostate cancer cells. Mol Cancer Ther 14, 586–596.

Dodd, K. M., Yang, J., Shen, M. H., Sampson, J. R., and Tee, A. R. (2015). mTORC1 drives HIF-1alpha and VEGF-A signalling via multiple mechanisms involving 4E-BP1, S6K1 and STAT3. Oncogene 34, 2239–2250.

Dowling, R. J., Lam, S., Bassi, C., Mouaaz, S., Aman, A., Kiyota, T., Al-Awar, R., Goodwin, P. J., and Stambolic, V. (2016). Metformin Pharmacokinetics in Mouse Tumors: Implications for Human Therapy. Cell Metab 23, 567–568.

Dowling, R. J., Topisirovic, I., Alain, T., Bidinosti, M., Fonseca, B. D., Petroulakis, E., Wang, X., Larsson, O., Selvaraj, A., Liu, Y., et al. (2010). mTORC1-mediated cell proliferation, but not cell growth, controlled by the 4E-BPs. Science 328, 1172–1176.

Dowling, R. J., Zakikhani, M., Fantus, I. G., Pollak, M., and Sonenberg, N. (2007). Metformin inhibits mammalian target of rapamycin-dependent translation initiation in breast cancer cells. Cancer Res 67, 10804–10812.

Fan, J., Ye, J., Kamphorst, J. J., Shlomi, T., Thompson, C. B., and Rabinowitz, J. D. (2014). Quantitative flux analysis reveals folate-dependent NADPH production. Nature 510, 298-302.

Fendt, S. M., Bell, E. L., Keibler, M. A., Davidson, S. M., Wirth, G. J., Fiske, B., Mayers, J. R., Schwab, M., Bellinger, G., Csibi, A. *, et al.* (2013). Metformin decreases glucose oxidation and increases the dependency of prostate cancer cells on reductive glutamine metabolism. Cancer Res 73, 4429–4438.

Gameiro, P. A., Yang, J., Metelo, A. M., Perez-Carro, R., Baker, R., Wang, Z., Arreola, A., Rathmell, W. K., Olumi, A., Lopez-Larrubia, P., et al. (2013). In vivo HIF-mediated reductive carboxylation is regulated by citrate levels and sensitizes VHL-deficient cells to glutamine deprivation. Cell metabolism 17, 372–385.

Gandin, V., Masvidal, L., Hulea, L., Gravel, S. P., Cargnello, M., McLaughlan, S., Cai, Y., Balanathan, P., Morita, M., Rajakumar, A., et al. (2016). nanoCAGE reveals 5' UTR features that define specific modes of translation of functionally related MTOR-sensitive mRNAs. Genome Res 26, 636–648.

Gandin, V., Sikstrom, K., Alain, T., Morita, M., McLaughlan, S., Larsson, O., and Topisirovic, I. (2014). Polysome fractionation and analysis of mammalian translatomes on a genome-wide scale. Journal of visualized experiments : JoVE.

Gebhart, G., Gamez, C., Holmes, E., Robles, J., Garcia, C., Cortes, M., de Azambuja, E., Fauria, K., Van Dooren, V., Aktan, G., et al. (2013). 18F-FDG PET/CT for early prediction of response to neoadjuvant lapatinib, trastuzumab, and their combination in HER2-positive breast cancer: results from Neo-ALTTO. J Nucl Med 54, 1862–1868.

Geyer, C. E., Forster, J., Lindquist, D., Chan, S., Romieu, C. G., Pienkowski, T., Jagiello-Gruszfeld, A., Crown, J., Chan, A., Kaufman, B., et al. (2006). Lapatinib plus capecitabine for HER2-positive advanced breast cancer. N Engl J Med 355, 2733–2743.

Giard, D. J., Aaronson, S. A., Todaro, G. J., Arnstein, P., Kersey, J. H., Dosik, H., and Parks, W. P. (1973). In vitro cultivation of human tumors: establishment of cell lines derived from a series of solid tumors. J Natl Cancer Inst 51, 1417–1423.

Gingras, A. C., Gygi, S. P., Raught, B., Polakiewicz, R. D., Abraham, R. T., Hoekstra, M. F., Aebersold, R., and Sonenberg, N. (1999). Regulation of 4E-BP1 phosphorylation: a novel two-step mechanism. Genes & development 13, 1422–1437.

Gingras, A. C., Kennedy, S. G., O'Leary, M. A., Sonenberg, N., and Hay, N. (1998). 4E-BP1, a repressor of mRNA translation, is phosphorylated and inactivated by the Akt(PKB) signaling pathway. Genes & development 12, 502–513.

Gravel, S. P., Avizonis, D., and St-Pierre, J. (2016). Metabolomics Analyses of Cancer Cells in Controlled Microenvironments. Methods Mol Biol 1458, 273–290.

Gravel, S. P., Hulea, L., Toban, N., Birman, E., Blouin, M. J., Zakikhani, M., Zhao, Y., Topisirovic, I., St-Pierre, J., and Pollak, M. (2014). Serine deprivation enhances antineoplastic activity of biguanides. Cancer Res 74, 7521–7533.

Griss, T., Vincent, E. E., Egnatchik, R., Chen, J., Ma, E. H., Faubert, B., Viollet, B., DeBerardinis, R. J., and Jones, R. G. (2015). Metformin Antagonizes Cancer Cell Proliferation by Suppressing Mitochondrial-Dependent Biosynthesis. PLoS Biol 13, e1002309.

Grove, J. R., Banerjee, P., Balasubramanyam, A., Coffer, P. J., Price, D. J., Avruch, J., and Woodgett, J. R. (1991). Cloning and expression of two human p70 S6 kinase polypeptides differing only at their amino termini. Mol Cell Biol 11, 5541–5550.

Hanahan, D., and Weinberg, R. A. (2011). Hallmarks of cancer: the next generation. Cell 144, 646–674.

Hettmer, S., Schinzel, A. C., Tchessalova, D., Schneider, M., Parker, C. L., Bronson, R. T., Richards, N. G., Hahn, W. C., and Wagers, A. J. (2015). Functional genomic screening reveals asparagine dependence as a metabolic vulnerability in sarcoma. Elife 4.

Hill, J. M., Roberts, J., Loeb, E., Khan, A., MacLellan, A., and Hill, R. W. (1967). L-asparaginase therapy for leukemia and other malignant neoplasms. Remission in human leukemia. JAMA: the journal of the American Medical Association 202, 882–888.

Hsieh, A. C., Liu, Y., Edlind, M. P., Ingolia, N. T., Janes, M. R., Sher, A., Shi, E. Y., Stumpf, C. R., Christensen, C., Bonham, M. J., et al. (2012). The translational landscape of mTOR signalling steers cancer initiation and metastasis. Nature 485, 55–61.

Huang, X., Wullschleger, S., Shpiro, N., McGuire, V. A., Sakamoto, K., Woods, Y. L., McBurnie, W., Fleming, S., and Alessi, D. R. (2008). Important role of the LKB1-AMPK pathway in suppressing tumorigenesis in PTEN-deficient mice. Biochem J 412, 211–221.

Javeshghani, S., Zakikhani, M., Austin, S., Bazile, M., Blouin, M. J., Topisirovic, I., St-Pierre, J., and Pollak, M. N. (2012). Carbon source and myc expression influence the antiproliferative actions of metformin. Cancer Res 72, 6257–6267.

Jiang, L., Shestov, A. A., Swain, P., Yang, C., Parker, S. J., Wang, Q. A., Terada, L. S., Adams, N. D., McCabe, M. T., Pietrak, B., et al. (2016). Reductive carboxylation supports redox homeostasis during anchorage-independent growth. Nature 532, 255–258.

Jin, Q., Yuan, L. X., Boulbes, D., Baek, J. M., Wang, Y. N., Gomez-Cabello, D., Hawke, D. H., Yeung, S. C., Lee, M. H., Hortobagyi, G. N., et al. (2010). Fatty acid synthase phosphorylation: a novel therapeutic target in HER2-overexpressing breast cancer cells. Breast cancer research: BCR 12, R96.

Kalender, A., Selvaraj, A., Kim, S. Y., Gulati, P., Brule, S., Viollet, B., Kemp, B. E., Bardeesy, N., Dennis, P., Schlager, J. J., et al. (2010). Metformin, independent of AMPK, inhibits mTORC1 in a rag GTPase-dependent manner. Cell Metab 11, 390–401.

Kim, B. R., Yoon, K., Byun, H. J., Seo, S. H., Lee, S. H., and Rho, S. B. (2014). The anti-tumor activator sMEK1 and paclitaxel additively decrease expression of HIF-1alpha and VEGF via mTORC1-S6K/4E-BP-dependent signaling pathways. Oncotarget.

Kisfalvi, K., Eibl, G., Sinnett-Smith, J., and Rozengurt, E. (2009). Metformin disrupts crosstalk between G protein-coupled receptor and insulin receptor signaling systems and inhibits pancreatic cancer growth. Cancer Res 69, 6539–6545.

Ko, B. W., Han, J., Heo, J. Y., Jang, Y., Kim, S. J., Kim, J., Lee, M. J., Ryu, M. J., Song, I. C.,Jo, Y. S., and Kweon, G. R. (2016). Metabolic characterization of imatinib-resistant BCR-ABL T315I chronic myeloid leukemia cells indicates down-regulation of glycolytic pathway and low ROS production. Leuk Lymphoma 57, 2180–2188.

Komurov, K., Tseng, J. T., Muller, M., Seviour, E. G., Moss, T. J., Yang, L., Nagrath, D., and Ram, P. T. (2012). The glucose-deprivation network counteracts lapatinib-induced toxicity in resistant ErbB2-positive breast cancer cells. Mol Syst Biol 8, 596.

Kordes, S., Pollak, M. N., Zwinderman, A. H., Mathot, R. A., Weterman, M. J., Beeker, A., Punt, C. J., Richel, D. J., and Wilmink, J. W. (2015). Metformin in patients with advanced pancreatic cancer: a double-blind, randomised, placebo-controlled phase 2 trial. Lancet Oncol 16, 839–847.

Kozma, S. C., Ferrari, S., Bassand, P., Siegmann, M., Totty, N., and Thomas, G. (1990). Cloning of the mitogen-activated S6 kinase from rat liver reveals an enzyme of the second messenger subfamily. Proc Natl Acad Sci U S A 87, 7365–7369.

Krall, A. S., Xu, S., Graeber, T. G., Braas, D., and Christofk, H. R. (2016). Asparagine promotes cancer cell proliferation through use as an amino acid exchange factor. Nat Commun 7, 11457.

Lane, A. N., and Fan, T. W. (2015). Regulation of mammalian nucleotide metabolism and biosynthesis. Nucleic acids research 43, 2466–2485.

Larsson, O., Morita, M., Topisirovic, I., Alain, T., Blouin, M. J., Pollak, M., and Sonenberg, N. (2012). Distinct perturbation of the translatome by the antidiabetic drug metformin. Proc Natl Acad Sci U S A 109, 8977–8982.

Laughner, E., Taghavi, P., Chiles, K., Mahon, P. C., and Semenza, G. L. (2001). HER2 (neu) signaling increases the rate of hypoxia-inducible factor 1alpha (HIF-1alpha) synthesis: novel mechanism for HIF-1-mediated vascular endothelial growth factor expression. Mol Cell Biol 21, 3995–4004.

Lehninger, A. L., Nelson, D. L., and Cox, M. M. (2000). Lehninger principles of biochemistry, 3rd edn (New York: Worth Publishers).

Leprivier, G., Remke, M., Rotblat, B., Dubuc, A., Mateo, A. R., Kool, M., Agnihotri, S., El-Naggar, A., Yu, B., Somasekharan, S. P., et al. (2013). The eEF2 kinase confers resistance to nutrient deprivation by blocking translation elongation. Cell 153, 1064–1079.

Li, Y. M., Zhou, B. P., Deng, J., Pan, Y., Hay, N., and Hung, M. C. (2005). A hypoxia-independent hypoxia-inducible factor-1 activation pathway induced by phosphatidylinositol-3 kinase/Akt in HER2 overexpressing cells. Cancer Res 65, 3257–3263.

Ling, A. M., and Keech, D. B. (1966). Pyruvate carboxylase from sheep kidney. I. Purification and some properties of the enzyme. Enzymologia 30, 367–380.

Liu, X., Romero, I. L., Litchfield, L. M., Lengyel, E., and Locasale, J. W. (2016). Metformin Targets Central Carbon Metabolism and Reveals Mitochondrial Requirements in Human Cancers. Cell Metab 24, 728–739.

Locasale, J. W. (2013). Serine, glycine and one-carbon units: cancer metabolism in full circle. Nat Rev Cancer 13, 572–583.

Locasale, J. W., Grassian, A. R., Melman, T., Lyssiotis, C. A., Mattaini, K. R., Bass, A. J., Heffron, G., Metallo, C. M., Muranen, T., Sharfi, H., et al. (2011). Phosphoglycerate dehydrogenase diverts glycolytic flux and contributes to oncogenesis. Nature genetics 43, 869–874.

Lovly, C. M., and Shaw, A. T. (2014). Molecular pathways: resistance to kinase inhibitors and implications for therapeutic strategies. Clin Cancer Res 20, 2249–2256.

Lozzio, C. B., and Lozzio, B. B. (1975). Human chronic myelogenous leukemia cell-line with positive Philadelphia chromosome. Blood 45, 321–334.

Mattaini, K. R., Sullivan, M. R., and Vander Heiden, M. G. (2016). The importance of serine metabolism in cancer. J Cell Biol 214, 249–257.

Maxwell, P. H., Wiesener, M. S., Chang, G. W., Clifford, S. C., Vaux, E. C., Cockman, M. E., Wykoff, C. C., Pugh, C. W., Maher, E. R., and Ratcliffe, P. J. (1999). The tumour suppressor protein VHL targets hypoxia-inducible factors for oxygen-dependent proteolysis. Nature 399, 271–275.

Mayerhofer, M., Valent, P., Sperr, W. R., Griffin, J. D., and Sillaber, C. (2002). BCR/ABL induces expression of vascular endothelial growth factor and its transcriptional activator, hypoxia inducible factor-1alpha, through a pathway involving phosphoinositide 3-kinase and the mammalian target of rapamycin. Blood 100, 3767–3775.

McArthur, G. A., Puzanov, I., Amaravadi, R., Ribas, A., Chapman, P., Kim, K. B., Sosman, J. A., Lee, R. J., Nolop, K., Flaherty, K. T., et al. (2012). Marked, homogeneous, and early [18F]fluorodeoxyglucose-positron emission tomography responses to vemurafenib in BRAF-mutant advanced melanoma. J Clin Oncol 30, 1628–1634.

Metallo, C. M., Gameiro, P. A., Bell, E. L., Mattaini, K. R., Yang, J., Hiller, K., Jewell, C. M., Johnson, Z. R., Irvine, D. J., Guarente, L., et al. (2011). Reductive glutamine metabolism by IDH1 mediates lipogenesis under hypoxia. Nature 481, 380–384.

Miloslavski, R., Cohen, E., Avraham, A., Iluz, Y., Hayouka, Z., Kasir, J., Mudhasani, R., Jones, S. N., Cybulski, N., Ruegg, M. A., et al. (2014). Oxygen sufficiency controls TOP mRNA translation via the TSC-Rheb-mTOR pathway in a 4E-BP-independent manner. Journal of molecular cell biology 6, 255–266.

Morita, M., Gravel, S. P., Chenard, V., Sikstrom, K., Zheng, L., Alain, T., Gandin, V., Avizonis, D., Arguello, M., Zakaria, C., et al. (2013). mTORC1 controls mitochondrial activity and biogenesis through 4E-BP-dependent translational regulation. Cell metabolism 18, 698–711.

Morita, M., Gravel, S. P., Hulea, L., Larsson, O., Pollak, M., St-Pierre, J., and Topisirovic, I. (2015). mTOR coordinates protein synthesis, mitochondrial activity and proliferation. Cell Cycle 14, 473–480.

Mullen, A. R., Wheaton, W. W., Jin, E. S., Chen, P. H., Sullivan, L. B., Cheng, T., Yang, Y., Linehan, W. M., Chandel, N. S., and DeBerardinis, R. J. (2012). Reductive carboxylation supports growth in tumour cells with defective mitochondria. Nature 481, 385–388.

Pause, A., Belsham, G. J., Gingras, A. C., Donze, O., Lin, T. A., Lawrence, J. C., Jr., and Sonenberg, N. (1994). Insulin-dependent stimulation of protein synthesis by phosphorylation of a regulator of 5'-cap function. Nature 371, 762–767.

Pavlova, N. N., and Thompson, C. B. (2016). The Emerging Hallmarks of Cancer Metabolism. Cell Metab 23, 27–47.

Phannasil, P., Ansari, I. H., El Azzouny, M., Longacre, M. J., Rattanapornsompong, K., Burant, C. F., MacDonald, M. J., and Jitrapakdee, S. (2017). Mass spectrometry analysis shows the biosynthetic pathways supported by pyruvate carboxylase in highly invasive breast cancer cells. Biochim Biophys Acta 1863, 537–551.

Pollak, M. (2013). Targeting oxidative phosphorylation: why, when, and how. Cancer Cell 23, 263–264.

Possemato, R., Marks, K. M., Shaul, Y. D., Pacold, M. E., Kim, D., Birsoy, K., Sethumadhavan, S., Woo, H. K., Jang, H. G., Jha, A. K., et al. (2011). Functional genomics reveal that the serine synthesis pathway is essential in breast cancer. Nature 476, 346–350.

Ravaud, A., Hawkins, R., Gardner, J. P., von der Maase, H., Zantl, N., Harper, P., Rolland, F., Audhuy, B., Machiels, J. P., Petavy, F., et al. (2008). Lapatinib versus hormone therapy in patients with advanced renal cell carcinoma: a randomized phase III clinical trial. J Clin Oncol 26, 2285–2291.

Ron D, H. H. C. S. H. L. P., Cold Spring Harbor, NY 2007). eIF2α phosphorylation in cellular stress-responses and disease. In Translational control in biology and medicine (Cold Spring Harbor, NY: Cold Spring Harbor Laboratory Press).

Roux, P. P., and Topisirovic, I. (2012). Regulation of mRNA translation by signaling pathways. Cold Spring Harbor perspectives in biology 4.

Sellers, K., Fox, M. P., Bousamra, M., 2nd, Slone, S. P., Higashi, R. M., Miller, D. M., Wang, Y., Yan, J., Yuneva, M. O., Deshpande, R., et al. (2015). Pyruvate carboxylase is critical for non-small-cell lung cancer proliferation. J Clin Invest 125, 687–698.

Semenza, G. L. (2007). Hypoxia-inducible factor 1 (HIF-1) pathway. Sci STKE 2007, cm8.

Semenza, G. L. (2013). HIF-1 mediates metabolic responses to intratumoral hypoxia and oncogenic mutations. J Clin Invest 123, 3664–3671.

Snell, K. (1984). Enzymes of serine metabolism in normal, developing and neoplastic rat tissues. Advances in enzyme regulation 22, 325–400.

Strumilo, S. (2005). Short-term regulation of the alpha-ketoglutarate dehydrogenase complex by energy-linked and some other effectors. Biochemistry (Mosc) 70, 726–729.

Sullivan, L. B., Gui, D. Y., Hosios, A. M., Bush, L. N., Freinkman, E., and Vander Heiden, M. G. (2015). Supporting Aspartate Biosynthesis Is an Essential Function of Respiration in Proliferating Cells. Cell 162, 552–563.

Sun, R. C., and Denko, N. C. (2014). Hypoxic regulation of glutamine metabolism through HIF1 and SIAH2 supports lipid synthesis that is necessary for tumor growth. Cell metabolism 19, 285-292

Tallarida, R. J. (2006). An overview of drug combination analysis with isobolograms. The of pharmacology and experimental therapeutics 319, 1–7

Thoreen, C. C., Chantranupong, L., Keys, H. R., Wang, T., Gray, N. S., and Sabatini, D. M. (2012). A unifying model for mTORC1-mediated regulation of mRNA translation. Nature 485, 109–113.

Trousil, S., Chen, S., Mu, C., Shaw, F. M., Yao, Z., Ran, Y., Shakuntala, T., Merghoub, T., Manstein, D., Rosen, N., et al. (2017). Phenformin enhances the efficacy of ERK inhibition in NF1-mutant melanoma. J Invest Dermatol.

Ursini-Siegel, J., Hardy, W. R., Zuo, D., Lam, S. H., Sanguin-Gendreau, V., Cardiff, R. D., Pawson, T., and Muller, W. J. (2008). ShcA signalling is essential for tumour progression in mouse models of human breast cancer. EMBO J 27, 910–920.

Ursini-Siegel, J., Rajput, A. B., Lu, H., Sanguin-Gendreau, V., Zuo, D., Papavasiliou, V., Lavoie, C., Turpin, J., Cianflone, K., Huntsman, D. G., and Muller, W. J. (2007). Elevated expression of DecR1 impairs ErbB2/Neu-induced mammary tumor development. Mol Cell Biol 27, 6361–6371.

Vander Heiden, M. G. (2011). Targeting cancer metabolism: a therapeutic window opens. Nature reviews Drug discovery 10, 671–684.

Vander Heiden, M. G., Cantley, L. C., and Thompson, C. B. (2009). Understanding the Warburg effect: the metabolic requirements of cell proliferation. Science 324, 1029–1033.

Vander Heiden, M. G., and DeBerardinis, R. J. (2017). Understanding the Intersections between Metabolism and Cancer Biology. Cell 168, 657–669.

Vazquez-Martin, A., Oliveras-Ferraros, C., Del Barco, S., Martin-Castillo, B., and Menendez, J.A. (2011). The anti-diabetic drug metformin suppresses self-renewal and proliferation of trastuzumab-resistant tumor-initiating breast cancer stem cells. Breast Cancer Res Treat 126, 355–364.

Walsh, A. J., Cook, R. S., Manning, H. C., Hicks, D. J., Lafontant, A., Arteaga, C. L., and Skala, M. C. (2013). Optical metabolic imaging identifies glycolytic levels, subtypes, and early-treatment response in breast cancer. Cancer research 73, 6164–6174.

Warner, J. R., Knopf, P. M., and Rich, A. (1963). A multiple ribosomal structure in protein synthesis. Proc Natl Acad Sci U S A 49, 122–129.

Wheaton, W. W., Weinberg, S. E., Hamanaka, R. B., Soberanes, S., Sullivan, L. B., Anso, E., Glasauer, A., Dufour, E., Mutlu, G. M., Budigner, G. S., and Chandel, N. S. (2014). Metformin inhibits mitochondrial complex I of cancer cells to reduce tumorigenesis. Elife 3, e02242.

Whelan, K. A., Schwab, L. P., Karakashev, S. V., Franchetti, L., Johannes, G. J., Seagroves, T. N., and Reginato, M. J. (2013). The oncogene HER2/neu (ERBB2) requires the hypoxia-inducible factor HIF-1 for mammary tumor growth and anoikis resistance. J Biol Chem 288, 15865–15877.

Wiernsperger, N. F., and Bailey, C. J. (1999). The antihyperglycaemic therapeutic and cellular mechanisms. Drugs 58 Suppl 1, 31-39; discussion 75-82.

Youngblood, V. M., Kim, L. C., Edwards, D. N., Hwang, Y., Santapuram, P. R., Stirdivant, S. M., Lu, P., Ye, F., Brantley-Sieders, D. M., and Chen, J. (2016). The Ephrin-A1/EPHA2 Signaling Axis Regulates Glutamine Metabolism in HER2-Positive Breast Cancer. Cancer research 76, 1825–1836.

Yuan, P., Ito, K., Perez-Lorenzo, R., Del Guzzo, C., Lee, J. H., Shen, C. H., Bosenberg, M. W., McMahon, M., Cantley, L. C., and Zheng, B. (2013). Phenformin enhances the therapeutic benefit of BRAF(V600E) inhibition in melanoma. Proc Natl Acad Sci U S A 110, 18226–18231.

Zakikhani, M., Dowling, R., Fantus, I. G., Sonenberg, N., and Pollak, M. (2006). Metformin is an AMP kinase-dependent growth inhibitor for breast cancer cells. Cancer Res 66, 10269–10273.

Zhang, B., Zheng, A., Hydbring, P., Ambroise, G., Ouchida, A. T., Goiny, M., Vakifahmetoglu-Norberg, H., and Norberg, E. (2017). PHGDH Defines a Metabolic Subtype in Lung Adenocarcinomas with Poor Prognosis. Cell Rep 19, 2289–2303.

Zhang, H., Li, H., Xi, H. S., and Li, S. (2012). HIF1alpha is required for survival maintenance of chronic myeloid leukemia stem cells. Blood 119, 2595–2607.

Zhang, J., Cao, J., Weng, Q., Wu, R., Yan, Y., Jing, H., Zhu, H., He, Q., and Yang, B. (2010). Suppression of hypoxia-inducible factor 1alpha (HIF-1alpha) by tirapazamine is dependent on eIF2alpha phosphorylation rather than the mTORC1/4E-BP1 pathway. PloS one 5, e13910

Zhang, J., Fan, J., Venneti, S., Cross, J. R., Takagi, T., Bhinder, B., Djaballah, H., Kanai, M., Cheng, E. H., Judkins, A. R., et al. (2014). Asparagine plays a critical role in regulating cellular adaptation to glutamine depletion. Mol Cell 56, 205–218.

Zhang, L., Han, J., Jackson, A. L., Clark, L. N., Kilgore, J., Guo, H., Livingston, N., Batchelor, K., Yin, Y., Gilliam, T. P., et al. (2016). NT1014, a novel biguanide, inhibits ovarian cancer growth in vitro and in vivo. J Hematol Oncol 9, 91.

Zhao, F., Mancuso, A., Bui, T. V., Tong, X., Gruber, J. J., Swider, C. R., Sanchez, P. V., Lum, J. J., Sayed, N., Melo, J. V., et al. (2010). Imatinib resistance associated with BCR-ABL upregulation is dependent on HIF-1alpha-induced metabolic reprograming. Oncogene 29, 2962–2972.

Zhao, Y., Liu, H., Liu, Z., Ding, Y., Ledoux, S. P., Wilson, G. L., Voellmy, R., Lin, Y., Lin, W., Nahta, R., et al. (2011). Overcoming trastuzumab resistance in breast cancer by targeting dysregulated glucose metabolism. Cancer Res 71, 4585–4597.

